# A role for myeloid miR-155 in regulating hypoxia induced seizures in neonatal C57BL/J6 mice

**DOI:** 10.1101/2022.09.22.508924

**Authors:** Devika Dahiya, Jonathan Smith, Tammy Strickland, Delphi Morris, Cristina Reschke, Tobias Engel, David Henshall, Claire E McCoy, Jennifer K Dowling

## Abstract

Hypoxic ischaemic injury (HIE) in the neonatal brain has significant consequences on neurodevelopment and increases the occurrence of neurological deficits in infants. HIE is also a leading cause of neonatal seizures. Therapeutic options for the treatment of HIE are very limited. Hypoxia-ischemia directly damages brain tissue in a primary-wave of injury which activates a cascade of events triggering local and systemic inflammatory responses, driven by the innate immune system, which contribute to a significant secondary-wave of injury taking place as early as 6 hours post-hypoxia-ischaemia. Levels of the well documented inflammatory microRNA, miR-155 are elevated in rodent seizure and epilepsy models. Here, we assessed the impact of, miR-155 deletion in myeloid cells, on regulating inflammation and seizure severity in a preclinical model of neonatal hypoxia-induced seizures (Hypoxia-Sz). Wildtype miR-155 (miR-155^+/+^LysMCre) mice were compared to a mouse line in which miR-155 was deleted in myeloid cells (miR-155^fl/fl^ LysMCre). We demonstrate significant upregulation of miR-155 target genes, brain-derived neurotrophic factor (*bdnf*), arginase-2 (*arg-2*), *ship-1* and *socs-1* in miR-155^fl/fl^ LysMCre mice compared to controls at various time points following Hypoxia-Sz. Conversely, we report decreased mRNA levels of pro-inflammatory cytokines IL-1β and IL-6 and lower protein levels of IL-1β in miR-155^fl/fl^ LysMCre mice as compared to WTs. Myeloid miR-155 deletion significantly reduced behavioural seizure severity score, reduced electrographically (EEG) measured seizure frequency and seizure burden as compared to mice to wildtypes, suggesting miR-155 regulation of seizure occurrence in this model. Behavioural tests for motor functions at 5 weeks post Hypoxia-Sz demonstrated differences between genotypes. Excitingly this work highlights that inhibition of miR-155, specifically in myeloid cells, may hold therapeutic benefit for both seizures and comorbidities associated with hypoxic brain injury.

## Background

Hypoxic Ischemic Encephalopathy (HIE) denotes neurologic dysfunction that occurs when the brain does not receive enough oxygen. The WHO identified birth asphyxia in the 5-leading causes of death amongst children under 5 years of age (1). Permanent motor-disorder cerebral palsy (CP) is associated with localised hypoxic injury (HI) (2, 3). Adults with CP have a 75% increased risk of non-communicable diseases, including asthma (4), depression and anxiety (5). Therapeutic hypothermia (TH) remains the only care presently used to reduce death and disability, yet is effective in just a select group of infants (6, 7).

On the other hand, global hypoxia in the brain triggers seizures in both rodent-models and humans, with infants becoming prone to epileptogenic tendencies (8–10). Hypoxia is in fact the most common cause of seizures in the first 24 hours of life (11). There are limited treatment options for hypoxia-induced seizures in infants. Successful treatment with anti-epileptic drugs (AEDs) including phenobarbital is poor. Additionally, this approach does not protect the immature brain from the neuroinflammation associated with hypoxic injury or decrease the risk of long-term neurological comorbidities (12). Thus, there is a major unmet clinical need for new therapeutic approaches in the treatment of hypoxic brain injury in infants.

Hypoxia-ischemia directly damages brain tissue in a primary-wave of injury caused by an energy failure occurring during periods of low aerobic respiration. The initial tissue injury activates a cascade of events triggering local and systemic inflammatory responses, which contribute to a significant secondary-wave of injury (13–15), taking place as early as 6 hours post-HI. The innate immune system drives this response and is partially responsible for the injurious second-wave (13, 14, 16). MicroRNAs, single-stranded non-coding nucleic acids, have been identified as crucial regulators of innate immune function (17–20). One such master regulator, miR-155, interacts with numerous inflammatory pathways post-hypoxic tissue injury (21–23). Targeted regulation of miR-155 activity may provide a novel therapeutic avenue for modulating the inflammation that follows hypoxic ischemic insult in the neonatal brain.

Hypoxia is one cause of tissue injury that has been identified to potently induce miR-155 expression, classifying it as a hypoxamiR (21, 22, 24, 25). Rodent models of ischemic stroke demonstrated increased miR-155 expression in the brain and plasma and a protective effect of miR-155 inhibition following injury (26–30). *In vivo* inhibition of miR-155 following distal middle cerebral artery occlusion (dMCAO) in mice, supported cerebral vasculature and resulted in reduced infarct size 21 days post-injury (26). In comparison to controls, miR-155 inhibitor-injected mice demonstrated well preserved capillary tight junctions and microvascular integrity with consequently improved blood flow in the peri-infarct area. *In vivo* miR-155 inhibition following murine dMCAO also promoted functional recovery post-experimental stroke (27). Inhibitor-injected mice demonstrated increased expression of miR-155 target proteins SOCS-1, SHIP-1, and C/EBP-β with consequent decrease in pro-inflammatory CCL12 and CXCL3 cytokine expression at 7 days and significantly increased levels of major cytokines IL-10 and IL-4 (anti-inflammatory) IL-6, IL-5, and IL-17 (context-dependent dual anti-and pro-inflammatory) at 14 days after dMCAO.

Macrophages were the most abundant leukocyte infiltrating the brain in rodent models of focal cerebral ischemia (31). Prominent expression of chemokine monocyte chemoattractant protein-1 (MCP-1) was noted in cerebral ischemia in rats (32, 33). Macrophages recruited to hypoxic sites are found to have altered phenotypes and appear to be polarized towards a pro-inflammatory phenotype (34–39). Exposure of human macrophage cell lines to non-lethal hypoxia resulted in secretion of significant quantities of TNF, IL-1, IL-8 along with increased TNF receptor and intercellular adhesion molecule-1 (ICAM-1), demonstrating the tendency of macrophages to assume a pro-inflammatory phenotype in environments with a low oxygen tension. Focal cerebral ischemia triggers macrophage activation and microglial aggregation around the area of tissue damage, demarcating the infarct area, phagocytosing and clearing the debris over time (31). Importantly, miR-155 transforms the macrophage transcriptional signature; where induction results in pro-inflammatory polarisation and simultaneous anti-inflammatory suppression; predominantly via the downregulation of approx. 650 genes with inhibitory action on the immune response (40). miR-155 targets SHIP-1, downregulating its expression (41–51). Similarly, SOCS-1 is inhibited by miR-155 (43, 46, 47, 49, 51–55). SHIP-1 and SOCS-1 are negative regulators of Type 1 cytokines (including IFNs), TLRs and the PI3K/AKT pathway; thus miR-155 induced repression of these molecules culminates in upregulated expression of inflammatory mediators IL-1β, IL-6, IL-8, TNF and iNOS.

Importantly, miR-155 levels are elevated in rodent seizure and epilepsy models (56–60). There is a current gap in our understanding of the exact role for myeloid miR-155 following hypoxic brain injury in new-borns. Here we sought to investigate the specific pathophysiological changes mediated by deletion of myeloid miR-155 in a murine model of neonatal hypoxic seizures induced by global cerebral hypoxia (61). We made use of miR-155^fl/fl^LysMCre C57BL/J6 mice where miR-155 is selectively knocked down in cells of a myeloid lineage, including monocytes, macrophages and microglia. We evaluated the expression of known targets of miR-155 in response to hypoxic stimulus at varying time points post-injury. We also examined the effect of myeloid miR-155 on induction of epileptogenic activity in the CNS along with behavioural modifications in response to globalised hypoxia. As such, we evaluated electroencephalographic outputs following hypoxic insult and simultaneously measured behavioural seizures using a modified 5 point Morrison’s Scale, as reported previously (62).

We postulated that knockdown of myeloid miR-155 would attenuate inflammation and improve phenotypic outcomes in neonates with hypoxia-induced seizures. We show increased expression of negatively regulated miR-155 targets (e.g.: HIF-1α, SOCS-1, SHIP-1 and Arg2) in miR-155^fl/fl^ LysMCre pups in our experimental model as compared to controls, resulting in reduced expression of pro-inflammatory cytokines. We demonstrate knockdown of myeloid miR-155 reduced symptomatic seizure severity as measured by the Morrison scale and improved EEG outputs following hypoxia. Finally, we found improved motor function in miR-155^fl/fl^LysMCre mice who suffered hypoxic brain injury as neonates in comparison to wildtypes.

## Main Text

### Methods

#### Generation of miR-155fl/fl x LysMCre mouse strains

C57BL/6-Mir155tm1.1Ggard/J (miR-155 floxed) mice were crossbred with B6.129P2-Lyz2tm1(cre)Ifo/J (LysMCre) mice to generate an experimental (miR-155fl/fl x LysMCre) colony. In miR-155^fl/fl^ LysMCre mice, specific and efficient Cre–mediated deletion of the loxP– flanked miR-155 gene (non-coding B cell integration cluster (Bic) gene) (63) occurs selectively in myeloid cells. In brief, the breeding strategy used to generate miR-155^+/+^LysMCre and miR-155^fl/fl^LysMCre was as follows. Stage 1 involved breeding a miR-155^fl/fl^ C57BL/J6 mouse with a LysMCre C57BLJ/6 mouse to obtain heterozygous miR-155 floxed x LysMCre offspring. These heterozygotes were bred together in stage 2 to generate both homozygous and heterozygous miR-155 floxed x LysMCre mice. Stage 3 involved crossbreeding homozygotes to maintain experimental and control colonies used for experiments. Genotyping was performed to amplify floxed and LysMCre regions and confirm miR-155^+/+^LysMCre and miR-155^fl/fl^LysMCre groups. Herein referred to as wildtype (WT) and miR-155^fl/fl^LysMCre (KO), respectively. LysMCre primers: Forward, CCC AGA AAT GCC AGA TTA CG; Reverse, CTT GGG CTG CCA GAA TTT CTC and MiR-155 Tm1.1 Ggard/J primers: Forward TAA GGC GCA CTT CCT TTA CA; Reverse ACG TAC ATC CCA CAG GTC ACT.

### Mouse Model of Neonatal Hypoxia-Induced Seizures

All procedures in the study including globalised hypoxia were performed in accordance to the European Communities Council Directives (86/609/EU and 2010/63/EU) and were reviewed and approved by the Research Ethics Committee of the Royal College of Surgeons in Ireland (REC #202002008) under license from the Health Products Regulatory Authority in Ireland (Global hypoxia model – AE19127/P013). C57BL/J6 miR-155^fl/fl^LysMCre (KO) and C57BL/J6 miR-155^+/+^LysMCre (WT) mice were sourced from the Biomedical Research Facility (BRF) at RCSI. Mating pair and trios were housed in a barrier-controlled facility within the Microsurgical Research and Training Facility (MRTF) on a 12-hour light-dark cycle with access to food and water ad libitum. Litters of male and female C57BL/J6 miR-155^fl/fl^LysMCre (KO) and C57BL/J6 miR-155^+/+^LysMCre (WT) mice were kept with their dams.

Neonatal mice [weight, 4–6g; age, postnatal day 6.5–7.5 (P7)] from the same litter were randomly assigned to either the normoxic control group or the hypoxic experimental group. Procedures for inducing hypoxia in mouse pups were based upon techniques previously described (61). P7 pups were exposed to decreasing flow rates of N2 across 8 minutes in a chamber until 5% O2/95% N2 were reached to produce globalised hypoxia. N2 flow rates were gradually reduced during the last 2 minutes of hypoxia to prevent ear drum damage in the pups. Oxygen levels were measured using a fibre optic oxygen meter to confirm efficacy of the protocol used **(supplemental figure 1)**. Non-hypoxic control pups were placed in the chamber but exposed to 21% O2 for an equivalent amount of time. The chamber was placed in an incubator and temperature controlled at 34°C and humidity at 80%.

**Figure 1.**
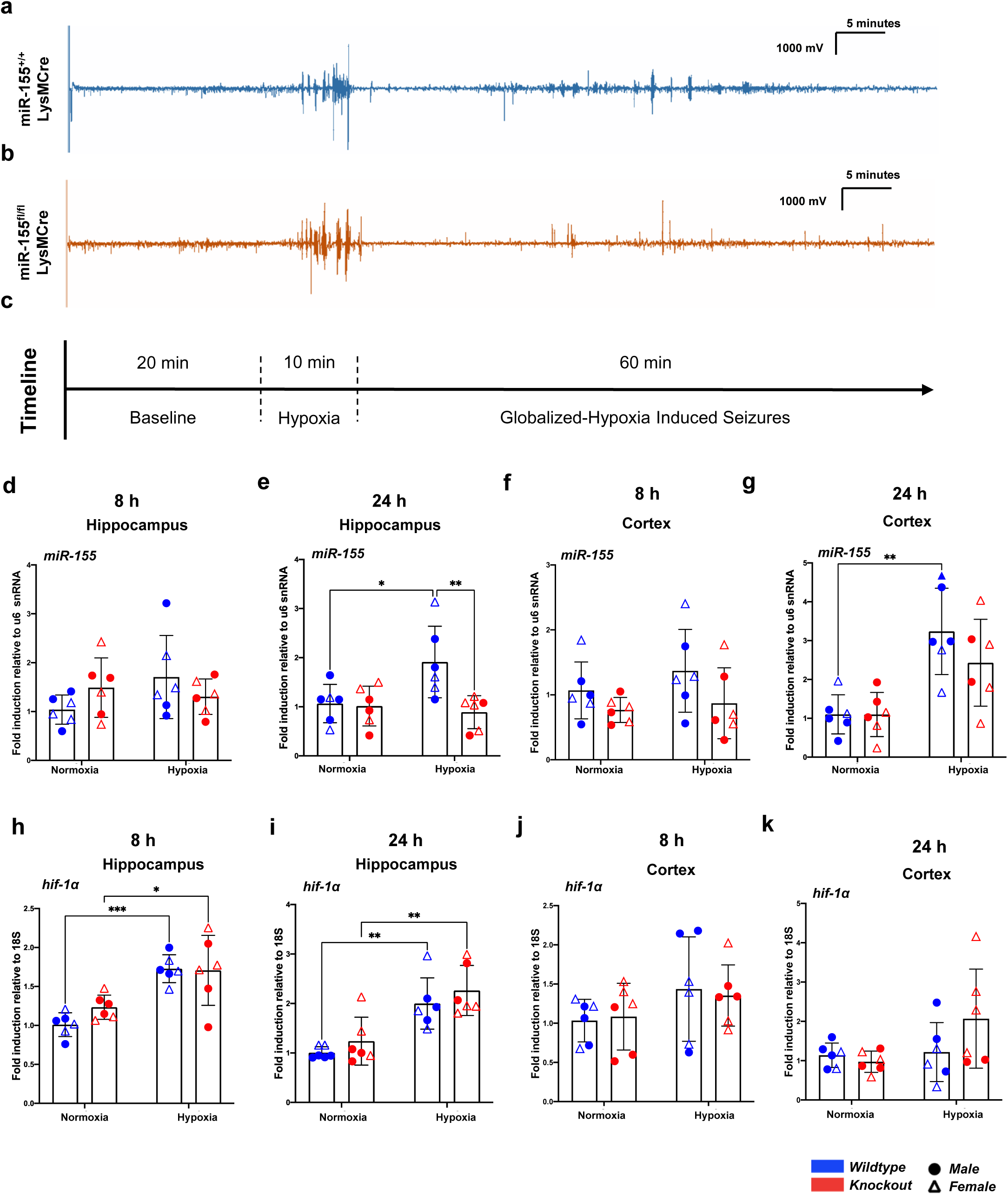
Induction of electroencephalographic seizures, miR-155 and HIF-1α in neonatal mice in response to globalised hypoxia. **(a-k)** WT (miR-155^+/+^LysMCre and **(b)** KO (miR-155^fl/fl^LysMCre) P7 pups were exposed to normoxia (21% oxygen) or hypoxia (5% oxygen) for 10 minutes at 34°C. Pups were monitored for 1 hour after hypoxia. **(a** and **b)** Representative electroencephalographic (EEG) trace of baseline recordings, hypoxic induction of seizure phenotype (10 min, 5% oxygen) and seizures 1 hour post hypoxic injury in **(a)** WT (miR-155^+/+^LysMCre and **(b)** KO (miR-155^fl/fl^LysMCre) neonatal mice. **(a) (c)** Experimental timeline, indicating baseline EEG recording, globalized hypoxic induction of the seizure phenotype, and seizures following hypoxic brain injury. **(d-k)** *miR-155* and *hif-1α* gene expression was compared in WT and miR-155^fl/fl^LysMCre P7 mice after hypoxic injury. Cortex and hippocampal tissue was extracted 8, 24 or 72 hours post procedure. RNA was extracted, cDNA generated and *miR-155* or *hif-1α* mRNA levels evaluated by RT-PCR. Hippocampal *miR-155* mRNA levels in P7 mice **(d)** 8 hours and **(e)** 24 hours after hypoxia. Cortical *miR-155* mRNA levels in P7 mice **(f)** 8 hours and **(g)** 24 hours after hypoxia. Hippocampal *hif-1α* mRNA levels in p7 mice **(h)** 8 hours and **(i)** 24 hours after hypoxia. Cortical *hif-1α* mRNA levels in P7 mice **(j)** 8 hours and **(k)** 24 hours after hypoxia. **(d-f)** *miR-155* expression was normalised to *u6* snRNA house-keeping gene and ΔCt of normoxic controls. **(h-k)** *hif-1α* mRNA was normalised to *18s* house-keeping gene and ΔCt of normoxic controls. Relative expression is mean ± SD of 6 biological replicates of each experimental category. * p˂0.05, ** p˂0.01, ***p˂0.001, two-way ANOVA. SHIP – Src Homology 2 domain-containing Inositol phosphatase 5-Phosphate, SOCS – Suppressor of Cytokine Signalling.

Mice pups were observed during and for 60 minutes after exposure to hypoxia. Behavioural seizures were scored using a 5 points Morrison scale modified for scoring hypoxia-induced seizures in neonatal mice (62) . Score 0: Normal behaviour; Score 1a: Immobility/motionless; Score 1b: Immobility and myoclonic jerks (spasms/shiver); Score 2: Rigid/ loss of posture; Score 3: Circling, swimming, peddling, tail extension; Score 4: Spasms, forelimb tonic-clonic; Score 5: Score 4 repeatedly. After hypoxia and 60 minutes of monitoring during the recovery period, the mouse pups were returned to the dam in the housing cages.

For electrographic recording, P7 mice pups were administered intraperitoneal buprenorphine hydrochloride 30 minutes before surgery (max 0.05 mg/kg). They were placed in a stereotaxic frame and anaesthetised with oxygen and isoflurane (5% for 1-2 min for induction, 2-3% for maintenance). A heating pad was used throughout the procedure to maintain body temperature and hindlimb pinch reflex was used throughout to assess the depth of anaesthesia regularly. Eye cream and appropriate eye protection was used to prevent the eyes drying out. Lidocaine/prilocaine (EMLA) cream and betadine was applied to the scalp. 3 partial craniotomy burr holes were performed and tethered electrodes (stainless steel screws soldered to a Teflon-insulated stainless steel wire; E/363/20, Bilaney Ltd., U.K. Diameter: 0.56 mm) were implanted to approximately 1 mm depth (avoiding the underlying brain surface) and secured into the skull with dental cement. One electrode was placed in each temporal lobe cortex and one reference in the cerebellum. Upon completion of the surgery, the mice pups were placed in an incubator at 34°C and humidity at 45% and monitored for 1 hour and 30 minutes while they recovered.

EEG outputs were taken at baseline for 20 minutes and then recorded during and for 1 hour after hypoxia using Xltek digital EEG-amplifier and digitalized with Twin software using Notch filter (1Hz high-pass and 60Hz low-pass) (Grass Technologies Ltd., Warwick, RI). Then, filtered files were uploaded to LabChart Pro (V7, ADInstruments Ltd.). Seizures were defined as electrographic polyspike discharges ≥5 Hz, ≥ 2x of baseline EEG amplitude and lasting ≥3 s. EEG total power [(μV2) is a function of EEG amplitude over time] was analyzed by integrating frequency bands from 0 to 60 Hz. Hypoxic ictal and inter-ictal traces were selected and the values were normalized to the pre-hypoxia baseline of each pup. The onset as well the number and duration of seizures (time from first spike to last spike) were calculated for each mouse pup. Total seizure burden was calculated as the cumulative time that pups were having electrographic polyspikes discharges.

### Mouse Tissue Samples

Pups were humanely euthanised at the end of the experiment via anaesthetic overdose. Pups were injected intraperitoneally with 0.05mg/kg of sodium pentobarbital. Following injection pups were assessed for pedal withdrawal response before tissue harvesting and sample collection was initiated. The pups were perfused with ice-cold saline to remove the intravascular blood components. The brains to be used for molecular and biochemical work were microdissected over wet ice into hippocampal and cortical samples which were then flash frozen in liquid nitrogen until being moved to -80°C for storage until tissue processing.

### Quantitative Real Time PCR

Hippocampi and Cortices were isolated and RNA was extracted using a TRIzol reagent. Samples were homogenized in 250 μl TRIzol reagent using non-reactive metal beads and a Tissue Lyser (Qiagen) for 4 minutes at 50 oscillations per second. RNA concentration was measured using a NanoDrop 100 spectrophotometer and RNA purity was measured by establishing the ratio of absorbance between 260 and 280 nm (260/280 ratio). Samples were normalised with RNAse-free water to 100 ng/μl or the lowest concentration, for use in Real-Time quantitative Polymerase Chain Reaction. A portion of the samples were made up to 10 ng/μl for miRNA RT-qPCR.

Quantitative real-time PCR was performed using a 7900HT Fast Real-Time PCR System (Applied Biosystems) in combination with High Capacity cDNA Reverse Transcription Kit (Applied Biosystems) or TaqMan MicroRNA Reverse Transcription Kit (Applied Biosystems) as per manufacturer’s protocol. Specific primers were purchased from Sigma Aldrich **(supplemental table 1)**.

### Cytokine Detection by Enzyme Linked Immunosorbant Assay (ELISA)

Samples were homogenized in LSLB **(supplemental table 2)** under the fume hood using non-reactive metal beads and a homogenizer machine. 250 μl of LSLB was added to the tissue along with 1 metal bead following which they were homogenised for 4 minutes at 50 oscillations per second. The samples were transferred to clean Eppendorf tubes for storage at -20°C.

After tissue homogenisation, the lysates and plasma samples were analysed for murine IL-1β using ELISA kits as per manufacturer’s instructions (R&D Biosystems). The plate was coated with capture antibody overnight. Samples were added in order to achieve a 1:2 dilution and incubated at 4°C overnight. Standards and a blank (reagent diluent) were added in technical replicates to generate a standard curve. Next, the detection antibody was added to the plate and incubated at room temperature for 2 hours. For visualisation of a signal, a working dilution of Streptavidin–HRP solution was added and incubated for 30 min at room temperature in a dark area. TMB substrate solution was added and plates were checked every 5 minutes for blue colour development. The enzymatic reaction was stopped by adding 2N sulfuric acid stop solution. The plate was read at 450nm absorbance peak using a spectrophotometer.

Linear regression analysis was conducted on Microsoft Excel to calculate IL-1β concentration in pg/ml. Final IL-1β concentrations were calculated as pg IL-1β/ μg of total brain protein using BCA assay results. Two-way ANOVA was conducted on PRISM (GraphPad Software Inc. California, USA) and used to determine statistical significance of differences between means of experimental groups.

### Cytometric Bead Array

Cytokine protein levels in brain lysates and plasma were measured using a BD™ CBA mouse inflammation kit. The CBA kit measures six proteins, IL-6, IL-10, TNF, IFN-γ, IL-12p70 and MCP-1 simultaneously by measuring the fluorescent intensities of bead populations coated with capture antibodies against each different cytokine. An aliquot of reconstituted standard was serially diluted 1:2 eleven times. Bead cocktails were made up and 50μl added to each sample and standard. PE-conjugated detection antibody was diluted as per the kit and 50μl was added to each sample and standard. Samples and Standards were incubated at room temperature in the dark for 2 hours. An acoustic focusing cytometer/FACS (Invitrogen) was used to measure fluorescent intensity following the recommended voltage settings for PE and APC-Cy7. Results were analysed using the FCAP Array™ Software and generated in a tabular and graphical format in Excel and PRISM (GraphPad Software Inc. California, USA). Two-way ANOVA was conducted on PRISM and used to determine statistical significance of differences between means of experimental groups.

## Motor Function

### Rotarod Test

The rotarod test was performed on P35 mice to assess gross motor function. The rotarod was set at a starting speed of 4rpm with an acceleration rate of 20 rpm/min. Mice were placed on the rotating rod facing away from the rotation direction by holding them by the tail. After 10 seconds from placing the mice on the rod, acceleration was started and the speed at which a 12mouse fell off was noted. If a mouse fell off before 10 seconds, the time of fall was noted and the mouse was returned to the rod up to three times. Falls before 5 seconds of placement were not recorded. The average time spent by the mouse on the rotating rod was then calculated. All analysis was carried out blinded to the experimental conditions. All files for the novel object recognition test were analysed as MP4 files through a media player and scored manually. Following unblinding, raw data from testing were grouped per treatment group and put through GraphPad Prism Software (Version 9; GraphPad Prism Inc., Chicago, III, USA) for statistical analysis.

## Results

### Globalised hypoxia induces seizure phenotype in miR-155^+/+^ and miR-155^fl/fl^ neonatal mice

We first confirmed the induction of sudden uncontrolled electrical brain activity in our model. 10 minutes of 5% oxygen in a temperature-controlled environment (34°C) induced seizure phenotype in P7 pups in both wildtype (miR-155^+/+^LysMCre) and knockout (miR-155^fl/fl^LysMCre) genotypes (**Fig 1A, B and C)**. As expected seizures were observed in the 1h recovery period at 21% oxygen following exposure to hypoxia, validating the globalised-hypoxic model of inducing seizure disease phenotype, as reported previously (61).

### HIF-1α and miR-155 expression levels are significantly elevated in neonatal mouse brain following exposure to hypoxia

Following Hypoxia-Sz, cortical and hippocampal brain tissue was harvested from mice at 8, 24 and 72 hours and gene expression evaluated. No significant differences were observed for miR-155 in the hippocampus or cortex 8 hours after hypoxic injury in WTs compared to normoxic controls (**Fig. 1D and F**). However, hypoxia significantly induced miR-155 expression in WT hippocampi (**Fig. 1E**) and cortices (**Fig. 1G**) 24 hours post hypoxia compared to the normoxia group. miR-155 was significantly reduced in hypoxia treated KO mice compared to WT mice in the hippocampus 24 hours after hypoxia (**Fig. 1E**). In contrast, cortical miR-155 levels were not significantly different in hypoxia treated KO compared to WT, suggesting hypoxic induction of miR-155 by non-myeloid cells within the cortex (**Fig. 1G**). miR-155 expression was not significantly altered in the hippocampus or the cortex of mice at 72 hours when comparing hypoxia treated groups to normoxia or when directly comparing WT and KO mice **(Supplementary Fig. 2).** Levels of transcription factor HIF-1α transcript were measured in order to characterise the hypoxic pathway upstream of miR-155, given miR-155 is driven by HIF1-α. mRNA levels of HIF1-α were significantly higher in hypoxia treated WT and KO hippocampi at 8 and 24 hours compared to normoxic controls (**Fig. 1H and I**). Cortical HIF-1α transcript levels trended upwards at 8 and 24 hours in both WT and KO mice, however this was not statistically significant (**Fig. 1J and K**). HIF-1α expression was not significantly changed 72 hours after hypoxia in the hippocampus or the cortex of mice when comparing hypoxia treated groups to normoxia controls or when directly comparing WT and KO mice (**Supplementary Fig. 2**). HIF-1α transcript expression in both WT and KO mice following Hypoxic-Sz demonstrates stepwise hypoxic induction of the transcription factor, which in turn affects miR-155 levels.

### Increased expression of negative regulators of inflammation in miR-155^fl/fl^ LysMCre neonates after Hypoxia-Sz

mRNA levels of Src homology-2 domain containing inositol 5-phosphate-1 (SHIP-1), a protein with anti-inflammatory actions and a target of miR-155, was assessed in WT and KO mice following Hypoxia-Sz. There was no change in *ship-1* mRNA levels at any timepoint in WT mice either in the hippocampus or cortex when comparing hypoxia group to normoxic controls (**Fig. 2A-C** and **supplemental Fig. 3**). However, *ship-1* was significantly upregulated at 8 and 24 hours after hypoxia in the hippocampus of KO mice (**Fig. 2A** and **B**). A significant induction of *ship-1* was also noted 24 hours post-hypoxia in the cortex of KO mice compared to normoxic controls (**Fig. 2C**). Suppressor of Cytokine Signalling-1 (SOCS-1), another protein with anti-inflammatory action whose expression is typically downregulated by miR-155 was assessed by RT-PCR. Similar changes were noted with *socs-1* transcript as observed for *ship-1*. *Socs-1* mRNA levels were significantly induced 8 and 24 hours after hypoxia at the hippocampus in KO but not in wildtype mice (**Fig. 2D and E**). *Socs-1* was significantly higher in KO hippocampi at 8 and 24 hours after hypoxia. Cortical *socs-1* transcript was significantly higher 24 hours post-hypoxia at the cortex in KO mice but not WT (**Fig. 2F**). Levels of the transcript were significantly higher in KO mice than wildtype mice 24 hours after hypoxia in the cortex. These results demonstrate that deletion of miR-155 in myeloid cells has a negative regulatory effect on inflammation via enhanced expression of *ship-1* and *socs-1,* known regulators of cytokine signalling.

**Figure 2.**
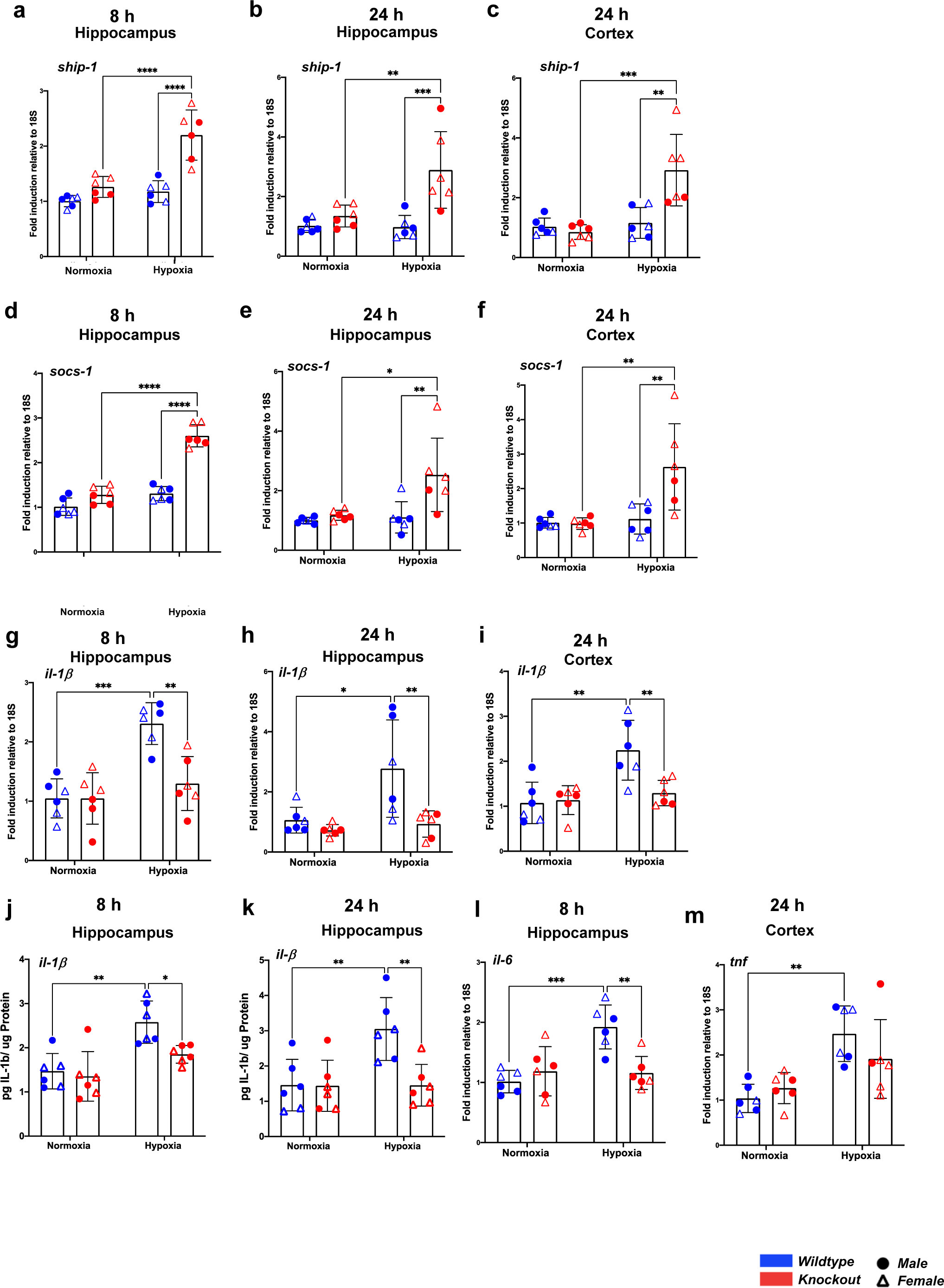
Myeloid miR-155 deletion reduces inflammation following hypoxic brain injury. **(a-m)** WT (miR-155^+/+^LysMCre) and myeloid KO (miR-155^fl/fl^ LysMCre) P7 mice underwent hypoxia/normoxia as in Fig. 1. Cortex and hippocampal tissue was extracted 8, 24 or 72 hours post procedure. RNA was extracted, cDNA generated and gene expression levels evaluated by RT-PCR. **(a-c)** Hippocampal *ship-1* mRNA levels in P7 mice at **(a)** 8 and **(b)** 24 hours. Cortical *ship-1* mRNA levels **(c)** 24 hours after hypoxia/normoxia. **(d-f)** Hippocampal *socs-1* mRNA levels in P7 mice at **(d)** 8 and **(e)** 24 hours. Cortical *socs-1* mRNA levels **(f)** 24 hours after hypoxia/normoxia. **(g-i)** Hippocampal *il-1β* mRNA levels in P7 mice at **(g)** 8 and **(h)** 24 hours. Cortical *il-1β* mRNA levels **(i)** 24 hours after hypoxia/normoxia. **(j,k)** Hippocampal *IL-1β* protein levels as measured by ELISA in P7 mice at **(j)** 8 and **(k)** 24 hours post hypoxic injury and expressed as picograms per micro gram of protein (pg/μg). **(a) (l)** Hippocampal *il-6* mRNA level in P7 mice at 8 hours following hypoxia. **(b) (m)** Cortical *tnf* mRNA level in P7l mice at 8 hours following hypoxia. (**a-i**) mRNA expression was normalised to *18s* house-keeping gene and ΔCt of normoxic controls. Fold change was calculated using the formula 2-(ΔCt). Relative expression is mean ± SD of 6 biological replicates of each experimental category. * p˂0.05, ** p˂0.01, ***p˂0.001, two-way ANOVA. pg – pictograms. SHIP – Src Homology 2 domain-containing Inositol phosphatase 5-Phosphate, SOCS – Suppressor of Cytokine Signalling, TNF – Tumour Necrosis Factor.

**Fig 3.**
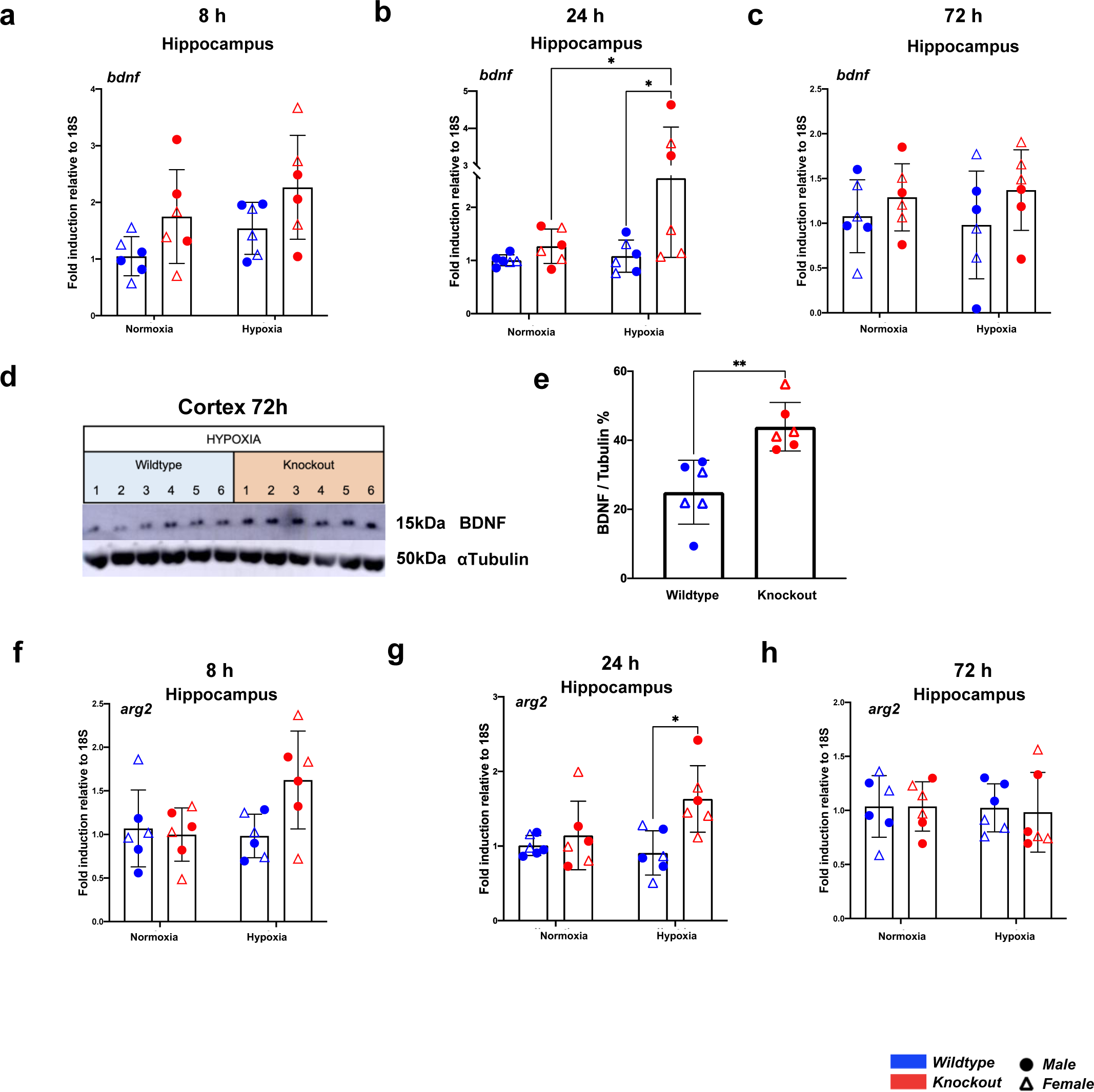
Myeloid miR-155 deletion increases transcriptional expression of inflammatory regulators BDNF and ARG-2 in response to hypoxic brain injury. **(a-g)** WT and myeloid KO P7 mice underwent hypoxia/normoxia as in Fig. 1. Cortex and hippocampal tissue was extracted 8, 24 or 72 hours after hypoxia/normoxia. Tissue was processed for RT-PCR and Western Blotting. For RT-PCR, RNA was extracted, cDNA generated and gene expression levels evaluated by RT-PCR. For western blotting tissue was added to low stringency lysis buffer (LSLB) and homogenized. Protein levels were was normalised by BCA assay (PIERCE). **(a - c)** Hippocampal *bdnf* mRNA levels in P7 mice at (a) 8 hours (b) 24 hours and (c) 72 hours following hypoxia/normoxia. **(a) (d)** Representative western blot of cortical BDNF protein levels in neonatal mice with hypoxic brain injury 72 hours after hypoxic injury **(b) (e)** Densitometry analysis showing relative expression levels of BDNF (n=6) 72 hours after hypoxic injury **(f-h)** Hippocampal *arg2* mRNA levels levels in P7 mice at **(f)** 8 hours **(g)** 24 hours and **(h)** 72 hours following hypoxia/normoxia. mRNA expression was normalised to *18s* house-keeping gene and ΔCt of normoxic controls. Fold change was calculated using the formula 2-(ΔΔCt). Relative expression is mean ± SD of 6 biological replicates of each experimental category. * p˂0.05, ** p˂0.01, ***p˂0.001, two-way ANOVA. BDNF – Brain Derived Neurotrophic Factor, ARG – Arginase

### Hypoxic induction of seizures significantly increases IL-1β, IL-6 and TNF transcription in the brain

Given *ship-1* and *socs-1* transcription was significantly enhanced in the hippocampus and cortex of hypoxia treated KO mice we next sought to determine any transcriptional changes to pro-inflammatory cytokines in KOs after Hypozia-Sz as compared to WT mice. RT-PCR data showed significantly increased transcription of *il*-*1β*, 8 and 24 hours after hypoxic induction of seizures in hippocampi of WT mice (**Fig. 2G and H**). These changes were no longer present at 72 hours (**Supplementary Fig. 4C**). *il-1β* mRNA levels were also slightly higher in wildtype cortices 8 hours after Hypoxia-Sz, however this trend was insignificant (p value 0.1555) (**Supplementary Fig. 4D**) and reached significance at 24 hours (p value 0.0014) (**Fig. 2I**). No significant change was observed at 72 hours (**Supplementary Fig. 4F**). *Il-6* mRNA levels were significantly higher in wildtype mice 8 hours after hypoxia (**Fig. 2l**). Similar trends were seen at the cortex (**Supplementary Fig. 4J**) however this effect was not significant (p value 0.6127). *Il-6* mRNA levels trended upwards in WT mice 24 post hypoxia at the hippocampus and cortex, (p value 0.1376 and 0.5741, respectively) (**Supplementary Fig. 4H and K**). RT-PCR data showed significantly increased transcription of *tnf* only at the 24 hour in WT mice cortices (**Fig. 2M**). These results demonstrate upregulated neuroinflammation signalling pathways triggered by globalised hypoxia within the neonatal brain.

### Myeloid miR-155 deletion suppresses hypoxic induction of IL-6 and IL-1β

Our results demonstrate significant upregulation of *il-1β*, *il-6* and *tnf* transcripts in WT mice at various timepoints post hypoxia in the neonatal brain. Concurrently, we found that myeloid miR-155 deletion ameliorated *il*-6 mRNA expression induced by hypoxia as seen in WT mice hippocampi 8 hours post-hypoxia (**Fig. 2l**). Myeloid miR-155 knockdown also ameliorated induction of *il*-1β mRNA in the hippocampus 8 and 24 hours after hypoxia compared to WTs (**Fig. 2G and H**). Quantification of IL-1β protein in tissue homogenates showed that changes seen at the transcript level were reflected at the protein level, where IL-1β was significantly higher in WT hippocampi 8 hours and 24 hours after hypoxia as compared to normoxic controls (**Fig. 4J and K**). This increase in IL-1β protein levels was again ameliorated in KO mice hippocampi 8 and 24 hours following hypoxia.

**Fig 4.**
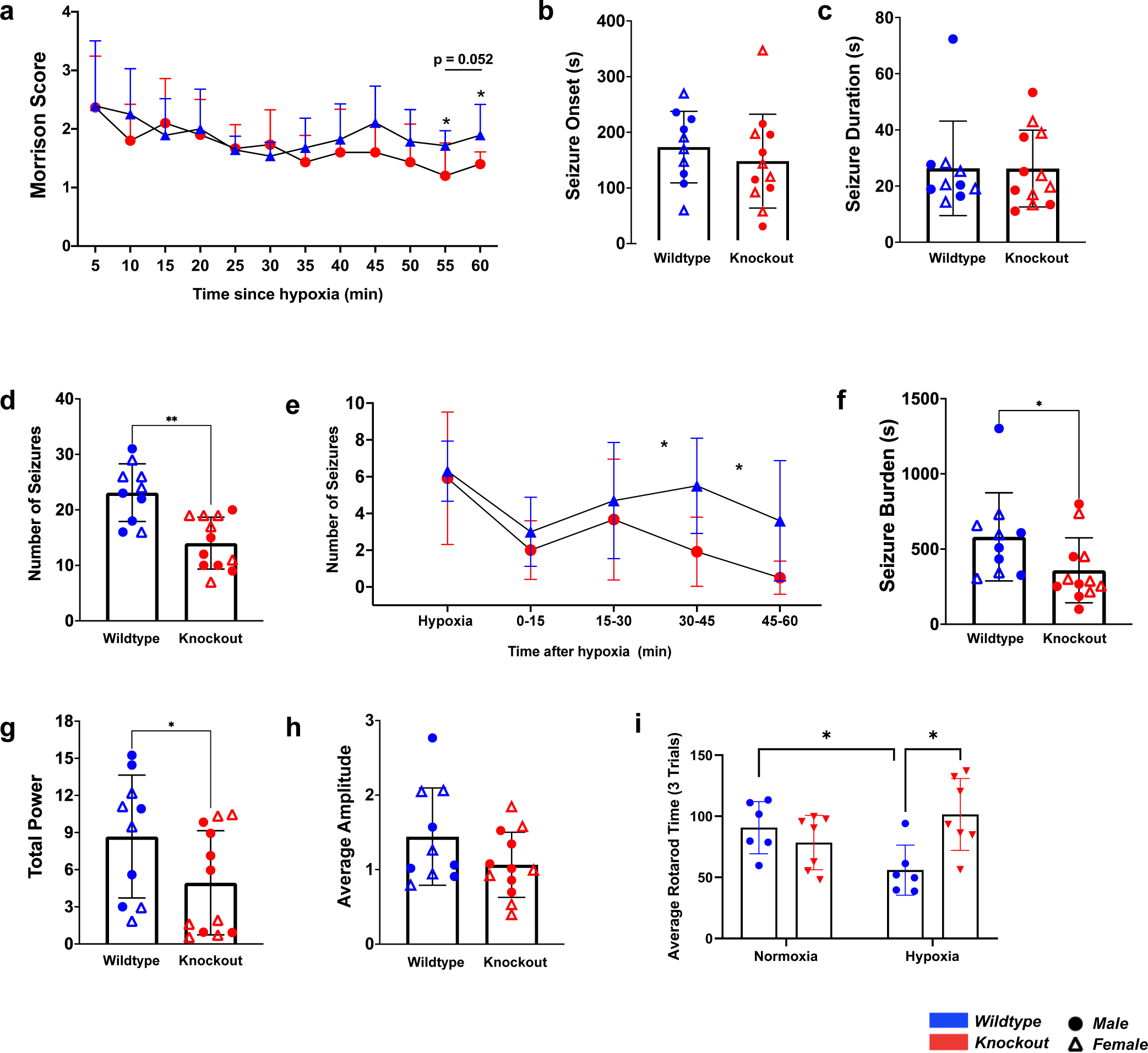
Myeloid miR-155 knockdown reduces seizure activity induced by hypoxic brain injury in the neonate. **(a)** Control and experimental genotype P7 pups were exposed to 10 min of hypoxia (5% oxygen) at 34° C following which behavioural seizures were monitored during a 1 hour recovery period. The highest Morrison scale seizure severity score reached every 5 minutes was noted and pooled from 14 WT and 15 miR-155^fl/fl^LysMCre mice. **(b – h)** Electrodes were surgically implanted in control and experimental genotype P7 pups. Baseline recordings (minimum 20 minutes) of resting brain activity was recorded. Mice were then exposed to hypoxia (5% oxygen) for 10 minutes at 34°C while continuously monitored using electroencephalography (EEG). Pups were monitored via EEG for 1 hour recovering after induction of seizures. **(a) (b)** Time to seizure onset in neonatal mice after initial exposure to hypoxia **(b)** Average seizure duration in neonatal mice **(c)** Seizure frequency during hypoxia and after hypoxic brain injury between wildtype and myeloid miR-155 knockout mice **(d)** The frequency of seizures in neonatal mice with time since hypoxic induction of seizure phenotype **(e)** Cumulative seizure burden in neonatal mice with hypoxia induced seizures **(f)** Total power of electroencephalographic outputs from neonatal mice following induction of seizure phenotype **(g)** Average amplitude of electroencephalographic outputs from neonatal mice following induction of seizure phenotype **(h)** Analysis of gross motor function in WT and KO mice following normoxia or hypoxia using the rotarod test (p-value 0.0167). Statistical analysis using 2way ANOVA tests (p-value <0.05). n=7-8 mice per group. Power and amplitude during hypoxia and recovery were normalised to baseline read outs. Average power and amplitude ± SD were calculated from 10 biological replicates of the control genotype and 12 biological replicates of the conditional knockout experimental category. * p˂0.05, ** p˂0.01, ***p˂0.001, unpaired nonparametric t-test (Mann-Whitney test) **(b-d and f-h**), multiple t-tests **(a,e)**.

Unfortunately, hippocampal and cortical TNF and IFNγ and hippocampal IL-6 were all below the limit of detection by cytometric bead array (CBA) (**Supplementary Fig. 5**). Protein levels of IL-10, IL-12p70 and MCP-1 were detectable in the neonatal brain by CBA, but no discernible pattern or trends were observed. Of note, we detected significantly reduced levels of the chemoattractant, MCP-1 in the periphery in plasma, 8 hours post HI in KO mice compared to WT mice **(Supplemental Fig. 7J)**. High levels of MCP-1 in human babies with HIE within the first 9 hours after birth was associated with death or severely abnormal neurodevelopment at 12 months of age (64). Collectively, our results demonstrate the importance of myeloid miR-155 in driving post-hypoxic inflammation.

### Knockdown of myeloid miR-155 increases inflammatory regulator Brain Derived Neurotrophic Factor (BDNF) mRNA and protein levels following Hypoxia-Sz

Brain Derived Neurotrophic Factor (BDNF) is a protein involved the physiological survival and growth of neurons. Dysregulation of BDNF has been identified in models of epilepsy, however, its role in hypoxia induced seizures is not well characterised. To investigate the miR-155-BDNF axis in our model of neuroinflammation, BDNF transcript and protein levels were examined. No changes in *bdnf* mRNA levels were detected in hypoxia treated WT mice at any timepoint compared to normoxic controls (**Fig. 3A-C**). However, mRNA levels of *bdnf* were significantly enhanced in hypoxia treated KO mice after 24 hours in the hippocampus compared to normoxia controls (**Fig. 3B**). Interestingly, hippocampal *bndf* mRNA levels appeared to be higher in KO mice under both normoxic and hypoxic conditions compared to WTs at the 8 hour and 72 hour timepoints, however this effect was not significant (**Fig. 3A** and **C**). While significant transcriptional changes in hippocampal *bdnf* transcript were observed at 24 hours post hypoxia, western blotting of BDNF protein across timepoints did not reach significance due to biological variability (**Supplementary Fig. 8**).

On the other hand, it was interesting that cortical *bdnf* mRNA expression trends seen at the transcriptional level while insignificant (**Supplementary Fig. 8**), translated into statistically significant protein expression changes at the cortex. Protein levels of BDNF were significantly higher 72 hours after hypoxia in KO mice compared to wildtype controls (**Fig. 3D** and **E**). BDNF protein levels 8 hours after hypoxia also mimicked changes at the transcriptional level where levels were slightly higher in KO mice than in WT mice, however this trend was insignificant (**Supplementary Fig. 9**). No differences were noted in cortical BDNF protein between KO and wildtype mice 24 hours after hypoxia, which matched transcriptional data (**Supplementary Fig. 8** and **9**).

### Arginase-2, a protein involved in metabolic reprogramming is enhanced in myeloid miR-155 knockout mice following Hypoxia-Sz

We have previously reported that arginase-2 (Arg-2) is a miR-155 regulated mitochondrial enzyme that plays an important role in regulating the metabolic reprogramming of macrophages from a pro-inflammatory phenotype towards an anti-inflammatory phenotype (65). Therefore we sought to examine the miR-155-Arg-2 axis in Hypoxic-Sz in wildtype and myeloid miR-155KO mice. RT-PCR data showed *arg-2* mRNA levels were significantly higher in KO mice than wildtypes 24 hours after induction of Hypoxic-Sz at the hippocampus (**Fig. 3G**). No significant changes were seen in terms of *arg-2* transcript expression at 8 or 72 hours after Hypoxic-Sz at the hippocampus, or at any timepoint examined post hypoxia in the cortex. While changes seen at the transcript level were not reflected at the protein level (**Supplemental Fig.9 D-F and J-L**), increased transcript levels of anti-inflammatory mediator *arg-2* in KO demonstrates that the miR-155-Arg-2 axis may be a means by which miR-155 inhibition has neuroprotective potential.

### Conditional knockout of myeloid miR-155 reduces hypoxia-induced behavioural seizure scores

A modified Morrison seizure scale was used to score visually apparent “behavioural” seizures in the pups following induction of the disease phenotype. More severe seizure characteristics resulted in a higher score. The highest score reached by a pup every 5 minutes was recorded for the 1-hour recovery period. Conditional knockout of myeloid miR-155 reduced the average seizure severity score reached by the pups over time during the 1-hour recovery period. WT and KO mice initially have similar seizure scores immediately after hypoxia, however the average seizure score graphs of the two genotypes separated with time **(Fig. 4A)**. The average seizure severity score was significantly lower in KO pups than in WT pups 55 and 60 minutes into the recovery period (p value 0.052) **(Fig. 4A)**.

### Myeloid miR-155 knock out significantly reduces seizure frequency, burden, power and improves motor function following globalised hypoxic injury in neontates

Seizure onset (duration into hypoxia seizures begin), duration (how long seizures last), frequency (the number of seizures during hypoxia and recovery) and burden (the cumulative time pups had seizures during hypoxia and recovery) were calculated and noted. Myeloid miR-155 deletion had no significant effect on seizure onset (148.2s vs 173.5s) or duration (26.2s vs 26.3s) as compared to WT mice (**Fig. 4B** and **C**). However, seizure frequency during hypoxia and recovery was significantly lower in KO mice compared to WTs (14.0 seizures vs 23.1 seizures) (**Fig. 4D**). A stratified analysis was conducted on seizure frequency to examine for changes over time, based on findings from Morrison seizure severity score findings. We found that KO and WT controls had similar seizure frequencies during hypoxia and during the first 15 minutes of the recovery (**Fig. 4E**). Seizure frequency in the KO group was slightly lower during the 15-30-minute period of recovery. The difference between the two genotypes increased during the 30-45 and 45-60 minute period of recovery, where the frequency of seizures was significantly lower in KO mice. Electroencephalographic (EEG) data also demonstrated that the overall seizure burden was lower in myeloid KO mice than wildtypes (358.7s vs 581.2s), and this effect was statistically significant (**Fig. 4F**).

Amplitude, (the deflection of the spike away from baseline (0 Hz) in both the positive and negative direction), and power, (the function of amplitude over time), were calculated from EEG data during the 1 hour recovery window for both genotypes. Average power was significantly lower in KO mice than wildtypes, (4.9 vs 8.7) (**Fig. 4G**). Similarly, average amplitude trended downwards in KO mice as compared to wildtypes, however this effect was not significant (1.1 vs 1.4) (**Fig. 4H**). Gross motor function was evaluated using the rotarod test at P35. There was no significant difference in motor function between WT and KO mice under normoxic conditions. There was a significant decrease in motor function in WT mice following hypoxia compared to normoxic controls. A significant improvement in motor function was observed in hypoxia treated KO mice compared to WTs **(Fig4. I).** These findings suggest that miR-155 dysregulation plays a significant role in the pathogenesis of Hypoxic-Sz *in vivo* and that miR-155 produced by myeloid cells majorly contributes to the disease phenotype and long term comorbidities.

## Discussion

Hypoxic ischaemic injury in the neonatal brain has significant consequences on neurodevelopment and increases the occurrence of neurological deficits in infants. Hypoxia-induced neonatal seizures are reported to cause lasting changes in brain excitability as well as cognitive and behavioural deficits due to an acute inflammatory response, with an increased risk of future conditions such as autism, epilepsy, and learning difficulties (66–68). Many existing medical interventions used to treat HIE work by reducing inflammation in the brain, however none specifically target hypoxia-ischaemia activated inflammatory molecular pathways (69–71).

Here we have addressed a knowledge gap, centred on the specific role for myeloid miR-155 in Hypoxia-Sz in neonates. The dysregulation of microRNAs has been reported in epilepsy extensively (72–74). In particular, aberrant activity of miR-155 demonstrably contributes to various models of epilepsy, and significantly, the inhibition or downregulation of miR-155 ameliorates epileptogenic activity across different models of epilepsy *in vivo* (56, 60, 75–78). To the best of our knowledge, we are the first to report significant upregulation of miR-155 in both the hippocampus and cortex of WT neonatal mice in Hypoxia-Sz.

We utilised miR-155^fl/fl^LysMCre C57BL/J6 mice where miR-155 is selectively knocked down in cells of a myeloid lineage. We used RT-PCR to assess the effect of myeloid miR-155 on regulating inflammation in the neonatal brain following Hypoxia-Sz. We measured expression of miR-155 targets and pro- and anti-inflammatory molecules in the brain from wildtype and miR-155^fl/fl^LysMCre neonatal mice after Hypoxic-Sz. We performed a stratified analysis, examining the hippocampus separately to the cortex instead of using whole brain lysates, due to increased vulnerability of the hippocampus to inflammation.

Our findings are consistent with previous reports of increases in pro-inflammatory cytokine expression in HIE (36, 62, 79, 80). We found Hypoxia-Sz was associated with increased expression of pro-inflammatory cytokines in WT neonatal mice. We also demonstrate that mRNA levels of miR-155 targets, *ship-1* and *socs-1*, known negative regulators of toll-like receptor (TLR) signalling and pro-inflammatory pathways, were significantly upregulated in miR-155^fl/fl^LysMCre mice following Hypoxia-Sz. This was associated with reduced proinflammatory cytokine IL-1β and IL-6 transcript expression as compared to wildtype control mice in the hippocampus. We also detected significant induction of IL-1β at both the transcriptional and protein level similar to a previous study using a neonatal mouse model of Hypoxia-Sz (62). This data suggests an anti-inflammatory and possible protective effect of myeloid deletion of miR-155 in Hypoxia-Sz. In contrast, our data did not reflect induction of TNF as seen in the other study. This may be explained by a refinement made in the model, where the duration of hypoxia was reduced in our hands from 15 to 10 minutes due to a higher mortality and attrition rates observed in our C57BL/J6 genotype compared to the C57BL/6OlaShD mice used in their study.

We also examined arginase-2, another target of miR-155 that we have previously reported to be crucial in the metabolic reprogramming of inflammatory macrophages to an anti-inflammatory phenotype (65, 81). Arg-2 dysregulation has been noted following hypoxic stimuli in the neonatal brain (82), and a deficiency of Arg-2 was shown to contribute to neuroinflammation and worsened pain-behaviours *in vivo* (83). We found increased transcription of *arg-2* in miR-155^fl/fl^LysMCre following Hypoxia-Sz as compared to WT controls. This result further validated our hypothesis that myeloid miR-155 is a key regulator of inflammation following hypoxic-induction of seizures *in vivo*. Certainly, other *in vivo* studies have demonstrated a reduction in proinflammatory pathways in total miR-155 knockout mice in the context of localised HI brain injury rather than globalised injury as in our study (30, 84, 85). However, the principals that miR-155 contributes to inflammation are comparable, especially when considering that miR-155 is highly expressed in macrophages (86, 87), and our study specifically examines the effect of myeloid miR-155 in hypoxic neuroinflammation.

It is important to highlight that while deletion of miR-155 in myeloid cells reduced the expression of inflammatory mediators, the seizure phenotype was not entirely resolved. Our data shows that deletion of myeloid miR-155 reduced disease severity *in vivo* as demonstrated by a significantly lower behavioural seizure severity score, seizure power, frequency and overall burden in miR-155^fl/fl^LysMCre as compared to WT controls. There was a significant relationship between miR-155 deletion and improvement in gross motor function following hypoxia in the rotarod test, further studies into the effects on fine motor function could be done using the balance beam test (63). Collectively, these findings suggest that miR-155 dysregulation plays a significant role in the pathogenesis of hypoxia-induced seizures *in vivo* and that miR-155 produced by myeloid cells majorly contributes to the disease phenotype.

Although seizure phenotype was not entirely resolved it is worth noting that the full behaviour of myeloid cells within the brain after globalised-hypoxic injury is not fully understood. Further characterisation of myeloid cell migration and activation in the brain and periphery is required to understand the full spectrum of effects myeloid miR-155 exerts in globalised hypoxic injury in the neonatal brain. Microglia also carry out important non-immune functions by interacting with components of the CNS embryonically and postnatally, contributing to its normal development (88, 89). As such, it is possible that knock-out of miR-155 in myeloid cells may result in structural and functional changes to components of the brain involved with the response to hypoxia. Additionally, microglia perform homeostatic functions including synaptic pruning, where faulty synapses are eliminated (88). However, synaptic pruning by microglia induces the development and aggravation of epilepsy by injurious stimuli due to the generation of an excitation/inhibition signal imbalance (90). As such, knock out of myeloid miR-155 may act to reduce the aggregative nature of pruning and consequently the development of seizure activity in the brain; while also exerting anti-inflammatory activity, and thus downregulating the development of epileptogenesis.

Importantly, we also examined for sex differences in this study. Rodent models of HIE have reported sex dependent differences in both mechanisms and outcomes of disease (91, 92). Additionally, differences in miR-155 target BDNF have been reported between males and females (93). We found no significant differences between males and females in any of our data. It is possible that the particular model of brain injury used in our study may hold limitations in being able to illicit and capture such differences, increasing our numbers may help draw out any differences between males and females in this population. Additionally, while examining our data, there appear to be outliers, however upon examining the data sets, the outlying data points were not from the same litter or experimental group. This suggests that what appears to be anomalous data may be accounted for by natural biological variation between animals, especially considering that these data points were generated from multiple litters and were in no instance all from one group of pups or tissues processed, reducing the likelihood that variations seen are due to experimental or human error. Additional experiments will help in better capturing the range of biological variation present and improve identification of true outliers.

miR-155 negatively regulates BDNF expression by targeted binding to the 3’UTR region of the molecules’ mRNA strand (94, 95). Direct interaction between miR-155 and BDNF was shown to mediate epileptogenic changes in the brain in murine temporal lobe epilepsy models (TLE) (96). Contrastingly, publications have reported a deleterious effect of BDNF in regulating seizure activity in the brain, linking high BDNF to adverse seizure outcomes (97, 98). While the exact role of BDNF in regulating seizures is not well characterised, we examined the myeloid-miR-155-BDNF axis in our model. Our data showed increased *bdnf* mRNA at 24 hours and protein expression at 72 hours in hypoxic treated miR-155^fl/fl^LysMCre brains compared to wildtype controls. Given the protective phenotype we have observed in miR-155^fl/fl^LysMCre mice post Hypoxia-Sz this result suggests that BDNF plays a neuroprotective role following globalised hypoxia induced seizures, and that myeloid miR-155 targets its downregulation post hypoxia.

Future work will look to better understand the effects that myeloid-miR-155 have on microglial function during neurodevelopment as well the activation and or temporal infiltration of microglia and macrophages following globalised hypoxic brain injury. It is also likely that any effects of myeloid miR-155 noted immediately following injury are from activated microglia which make up 1/10^th^ of the brain cellular population (99); and later effects mediated by infiltrating macrophages, however this axis requires further investigation. Differentiating between the specific functions microglial-miR-155 and macrophage-miR-155 hold in regulating inflammation following hypoxic injury will enable a more targeted approach to developing treatments. As such, immunomodulation, via the inhibition of myeloid produced miR-155, may serve to reduce seizure activity and consequently reduce neuronal injury, providing a novel therapeutic avenue for the management of neonatal hypoxic encephalopathy.

## Conclusions

We appreciate that our study holds certain limitations. While this model examines globalised brain injury following hypoxia, it does not mimic the more localised brain injury that occurs with mixed hypoxic-ischaemic insults in HIE. However, the information generated from this work can be used to guide our understanding of the role of myeloid miR-155 in HIE. While we observed reduced disease severity in miR-155^fl/fl^LysMCre pups we have not identified the exact sources of the miR-155. Our data indicates that miR-155 was certainly induced in the cortex of miR-155^fl/fl^LysMCre post hypoxia, suggesting an additional and non-myleoid source of miR-155 in the neonatal brain post hypoxia. However, its remains that myeloid deletion of miR-155 was sufficient to significantly reduce disease severity and seizure burden. Future work is required to elucidate the temporal and spatial source of miR-155 in this model and others models of HIE. Targeting inflammation by inhibition of miR-155 may protect the neonatal brain from injury via decreased expression of pro-inflammatory molecules at the transcriptional and protein level; reduced seizure severity, frequency and burden and improved motor functional outcomes later in life. From a clinical translation perspective, future studies should focus on the administration of an anti-miR-155 antagomiR in wildtype mice to verify the neuro-protective capacity of a targeted miR-155 approach in Hypoxic-Sz in the neonatal brain.

## Acknowledgments

This project has received funding from the European Union’s Horizon 2020 research and innovation programme under the Marie Skłodowska-Curie grant agreement No. 897880.

## Abbreviations

*ANOVA*: – Analysis of Variance
*Arg*: - Arginase
*ATP*: – Adenosine Triphosphate
*BBB*: – Blood Brain Barrier
*Bcl6*: – B-cell lymphoma 6 protein
*BDNF*: – Brain Derived Neurotrophic Factor
*Bic gene*: – B-cell integration cluster gene
*C/EBP*: – CCAAT/enhancer-binding proteins
*CBA*: – Cytometric Bead Array
*CNS*: – Central Nervous System
*CP*: – Cerebral Palsy
*DC*: – Dendritic Cell
*dMCAO*: – distal Middle Cerebral Artery Occlusion
*DMEM*: – Dulbecco’s Modified Eagle Medium
*DNA*: – Deoxyribonucleic Acid
*DSS*: – Dextran Sulphate Sodium
*EEG*: – Electroencephalograph
*ELISA*: – Enzyme Linked Immunosorbent Assay
*GC*: – Glucocorticoids
*HIE*: – Hypoxic Ischaemic Encephalopathy
*HIF*: – Hypoxia Inducible Factor
*HRI*: – Hypoxia Reperfusion Injury
*Hypoxia*-*Sz*/*Hypoxic*-*Sz*: – Hypoxia induced Seizures
*IFN*: – Interferon
*IL*: – Interleukin
*IP*: – Intraperitoneal
*IRF*: – Interferon Regulatory Factor
*IRI*: – Ischaemia Reperfusion Injury
*ISRE*: – Interferon-Sensitive Response Elements
*KSRP*: - KH-type Splicing Regulatory Protein
*LSLB*: – Low Stringency Lysis Buffer
*Maf*-*b*: – Musculoaponeurotic fibrosarcoma-oncogene homolog-bD
*MCP*: – Monocyte Chemoattractant Protein
*MMP*: – Matrix Metalloproteinase
*MRTF*: – Microsurgical Research and Training Facility
*NCS*: – Non-Convulsive Seizures
*NF*-*kB*: – Nuclear Factor kappa-light-chain-enhancer of activated B cells
*OGD*/*R*: – Oxygen Glucose Deprivation/Reoxygenation
*PBS*: – Phosphate Buffered Saline
*PI3K*-*AKT*: - Phosphatidylinositol-3-Kinase - Protein kinase B
*PRR*: – Pattern Recognition Receptors
*RCLB*: – Red Cell Lysis Buffer
*RISC*: – RNA Induced Silencing Complex
*RNA*: – Ribonucleic Acid
*ROS*: – Reactive Oxidative Species
*RT-PCR*: – Real Time quantitative Polymerase Chain Reaction
*SDS*-*PAGE*: – Sodium Dodecyl Sulphate–Polyacrylamide Gel Electrophoresis
*SHIP*-*1*: - Src homology 2 (SH2) domain containing inositol polyphosphate 5-phosphatase 1
*SOCS*-*1*: – Suppressor of Cytokine Signalling-1
*TBST*: - Tris-buffered Saline-Tween
*TF*: – Transcription Factor
*TGF*: – Transforming Growth Factor
*TH*: – Therapeutic Hypothermia
*TLE*: – Temporal Lobe Epilepsy
*TLR*: – Toll Like Receptor
*TNF*: – Tumour Necrosis Factor
*UTR*: – Untranslated Region
*WB*: – Western Blot
*WHO*: – World Health Organisation
*WT*: – Wildtype

## Supplementary Figure Legends

**Supplementary Figure. 1.**
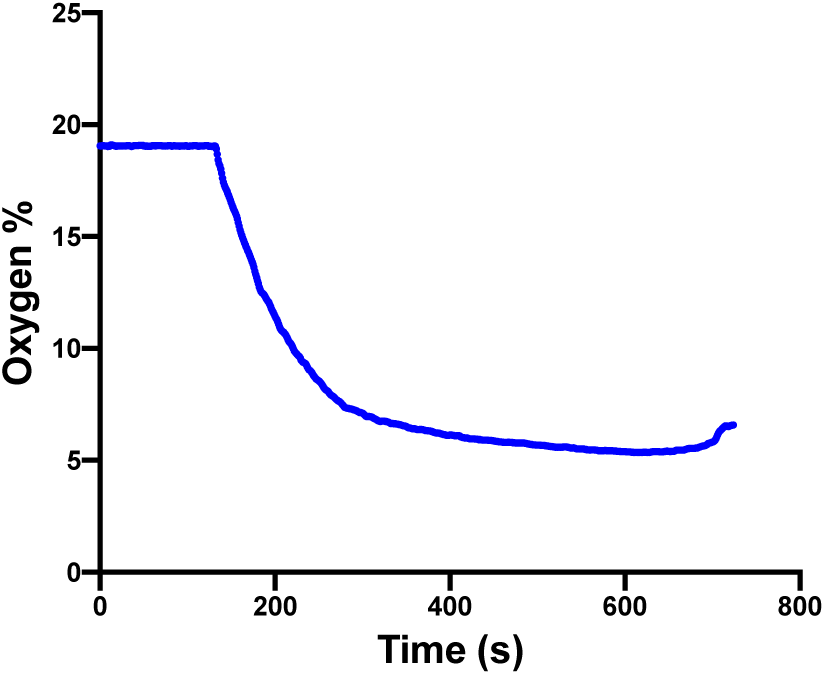
Oximeter recordings of oxgygen concentration. Changes in oxygen concentration from normal atmospheric levels to 5% O2/95% N2 with time were measured using a fibre optic oxygen meter (pyroscience).

**Table 1:**
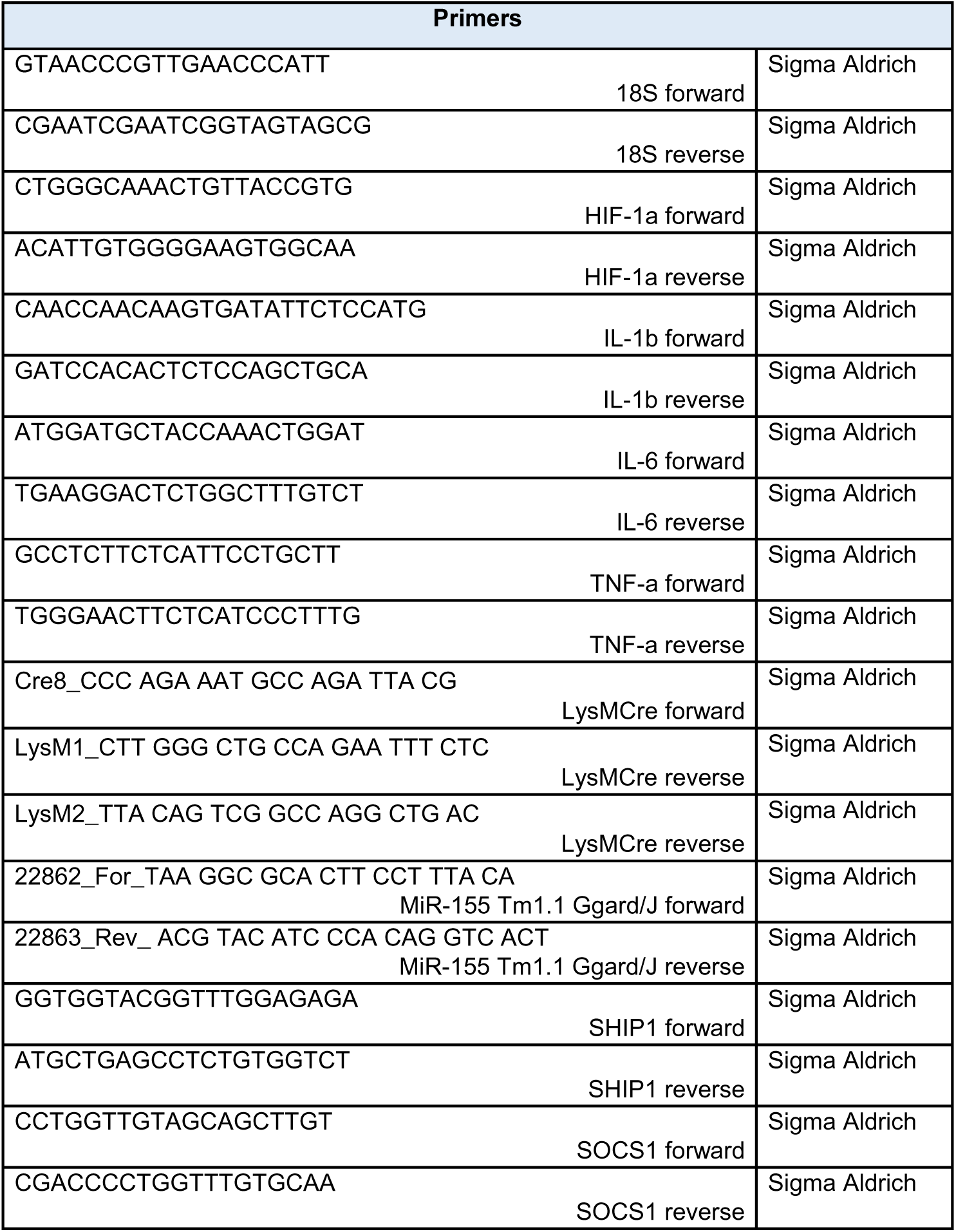
Primers, Antibodies & Commercial Kits.

**Table 2:** Reagents and concentrations used to make up LSLB, used to homogenize tissue samples for protein analysis.

|  |
| --- |
| Reagent |
| 50mM HEPES |
| 100mM NaCl |
| 10% Glycerol |
| 0.5% NP-40 |
| 1mM EDTA |
| dH<sub>2</sub>O |
| Protease inhibitor cocktail<br>x1/40ml |

**Supplemental Figure 2.**
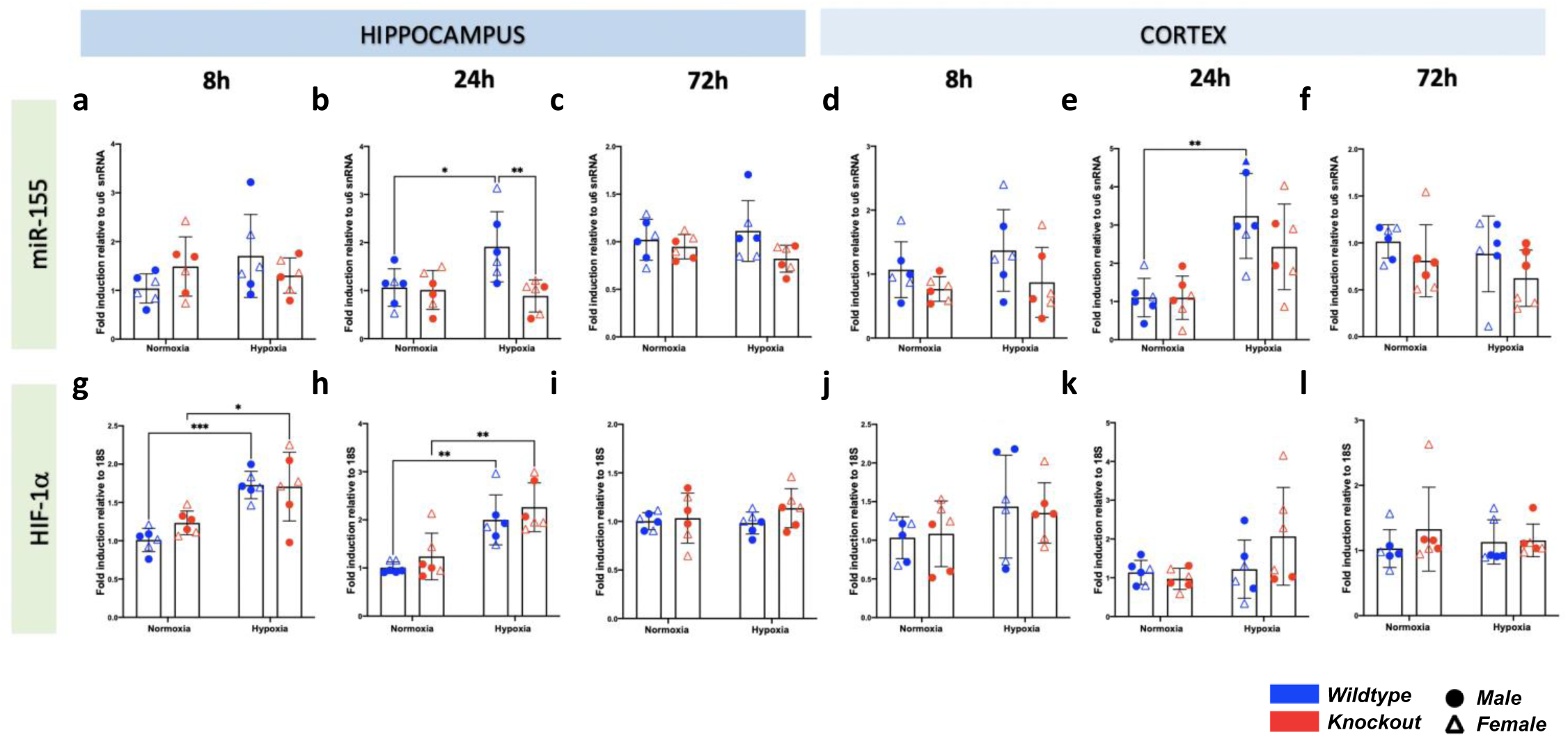
Induction of miR-155 miRNA and HIF-1α mRNA in neonatal mice in response to globalised hypoxia. Control and experimental genotype p7 pups were exposed to normoxia or hypoxia (5% oxygen) for 10 minutes at 34°C. Pups were monitored for 1 hour after induction of seizures. Brain tissue was extracted 8, 24 or 72 hours after induction of seizures by hypoxia. **(a-f)** Hippocampal and cortical miR-155 microRNA levels in neonatal mice with hypoxic brain injury at 8, 24 and 72 hours following induction of seizure phenotype **(g-l)** Hippocampal and cortical HIF-1α mRNA levels in neonatal mice with hypoxic brain injury at 8, 24 and 72 hours following induction of seizure phenotype RNA was extracted from tissue using TRIzol reagent protocols. cDNA was generated from samples and miR-155 or HIF-1α mRNA expression levels were evaluated using RT-qPCR. miR- 155 expression was normalised to u6 snRNA house-keeping gene and ΔCt of non-treated cells. HIF-1α mRNA expression was normalised to 18s house-keeping gene and ΔCt of non-treated cells. Fold change was calculated using the formula 2^-(ΔΔCt)^. Relative expression is mean ± SD of 6 biological replicates of each experimental category. * p˂0.05, ** p˂0.01, ***p˂0.001, two-way ANOVA. HIF – Hypoxia Inducible Factor

**Supplemental Fig 3.**
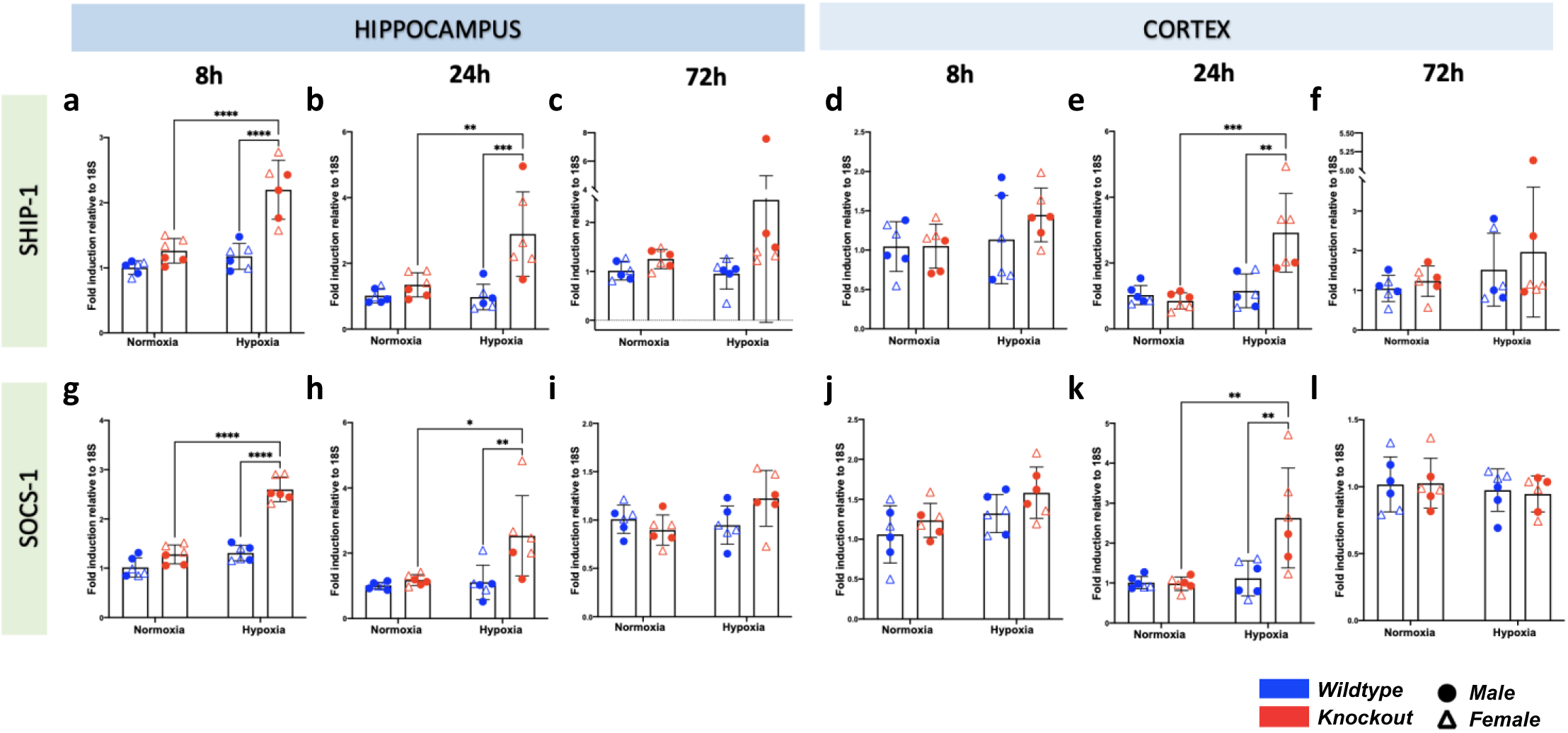
Increased transcription of anti-inflammatory proteins SHIP-1 and SOCS- 1 in neonatal mice in response to globalised hypoxia. Control and experimental genotype p7 pups were exposed to normoxia or hypoxia (5% oxygen) for 10 minutes at 34°C. Pups were monitored for 1 hour after induction of seizures. Brain tissue was extracted 8, 24 or 72 hours after induction of seizures by hypoxia. **(a-f)** Hippocampal and cortical *ship-1* mRNA levels in neonatal mice with hypoxic brain injury at 8, 24 and 72 hours following induction of seizure phenotype **(g-l)** Hippocampal and cortical *socs-1* mRNA levels in neonatal mice with hypoxic brain injury at 8, 24 and 72 hours following induction of seizure phenotype RNA was extracted from tissue using TRIzol reagent protocols. cDNA was generated from samples and *ship-1* or *socs-1* mRNA expression levels were evaluated using RT-qPCR. miR-155 expression was normalised to u6 snRNA house-keeping gene and DCt of non-treated cells. SOCS-1 mRNA expression was normalised to 18s house-keeping gene and DCt of non-treated cells. Fold change was calculated using the formula 2-(DDCt). Relative expression is mean ± SD of 6 biological replicates of each experimental category. * p˂0.05, ** p˂0.01, ***p˂0.001, two-way ANOVA. SHIP – Src Homology 2 domain-containing Inositol phosphatase 5-Phosphate, SOCS – Suppressor of Cytokine Signalling.

**Supplemental Figure 4.**
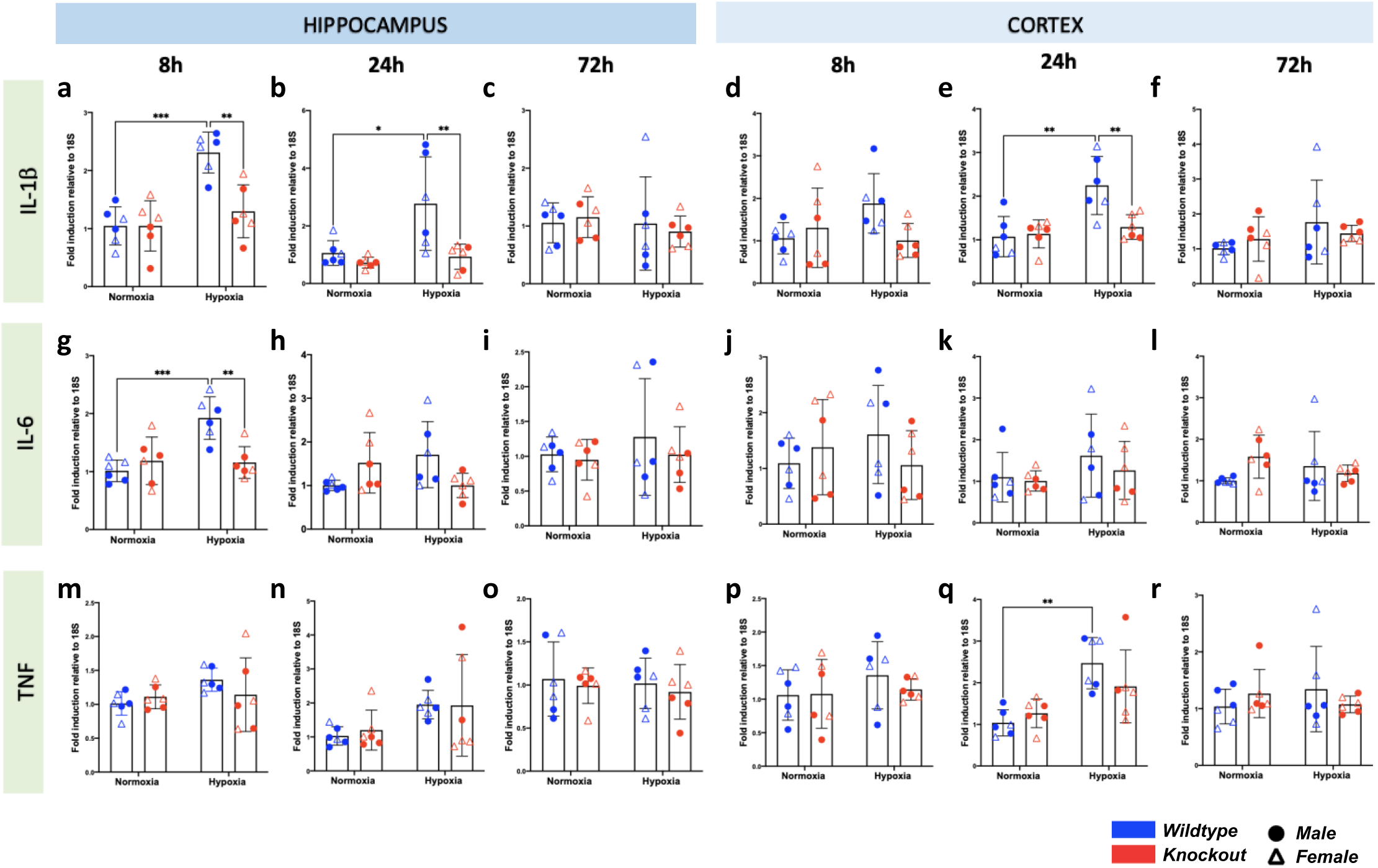
Increased transcription of inflammatory cytokines in neonatal mice in response to globalised hypoxia. Control and experimental genotype p7 pups were exposed to normoxia or hypoxia (5% oxygen) for 10 minutes at 34°C. Pups were monitored for 1 hour after induction of seizures. Brain tissue was extracted 8, 24 or 72 hours after induction of seizures by hypoxia. **(a-f)** Hippocampal and cortical *il-1β* mRNA levels in neonatal mice with hypoxic brain injury at 8, 24 and 72 hours following induction of seizure phenotype **(g-l)** Hippocampal and cortical *il-6* mRNA levels in neonatal mice with hypoxic brain injury at 8, 24 and 72 hours following induction of seizure phenotype **(m-r)** Hippocampal and cortical *tnf* mRNA levels in neonatal mice with hypoxic brain injury at 8, 24 and 72 hours following induction of seizure phenotype RNA was extracted from tissue using TRIzol reagent protocols. cDNA was generated from samples and *il-1β*, *il-6* or *tnf* mRNA expression levels were evaluated using RT-qPCR. miR-155 expression was normalised to u6 snRNA house-keeping gene and ΔCt of non-treated cells. Cytokine mRNA expression was normalised to *18s* house-keeping gene and ΔCt of non- treated cells. Fold change was calculated using the formula 2^-(ΔΔCt)^. Relative expression is mean ± SD of 6 biological replicates of each experimental category. * p˂0.05, ** p˂0.01, ***p˂0.001, two-way ANOVA. IL-Interleukin, TNF – Tumour Necrosis Factor.

**Supplemental Figure 5.**
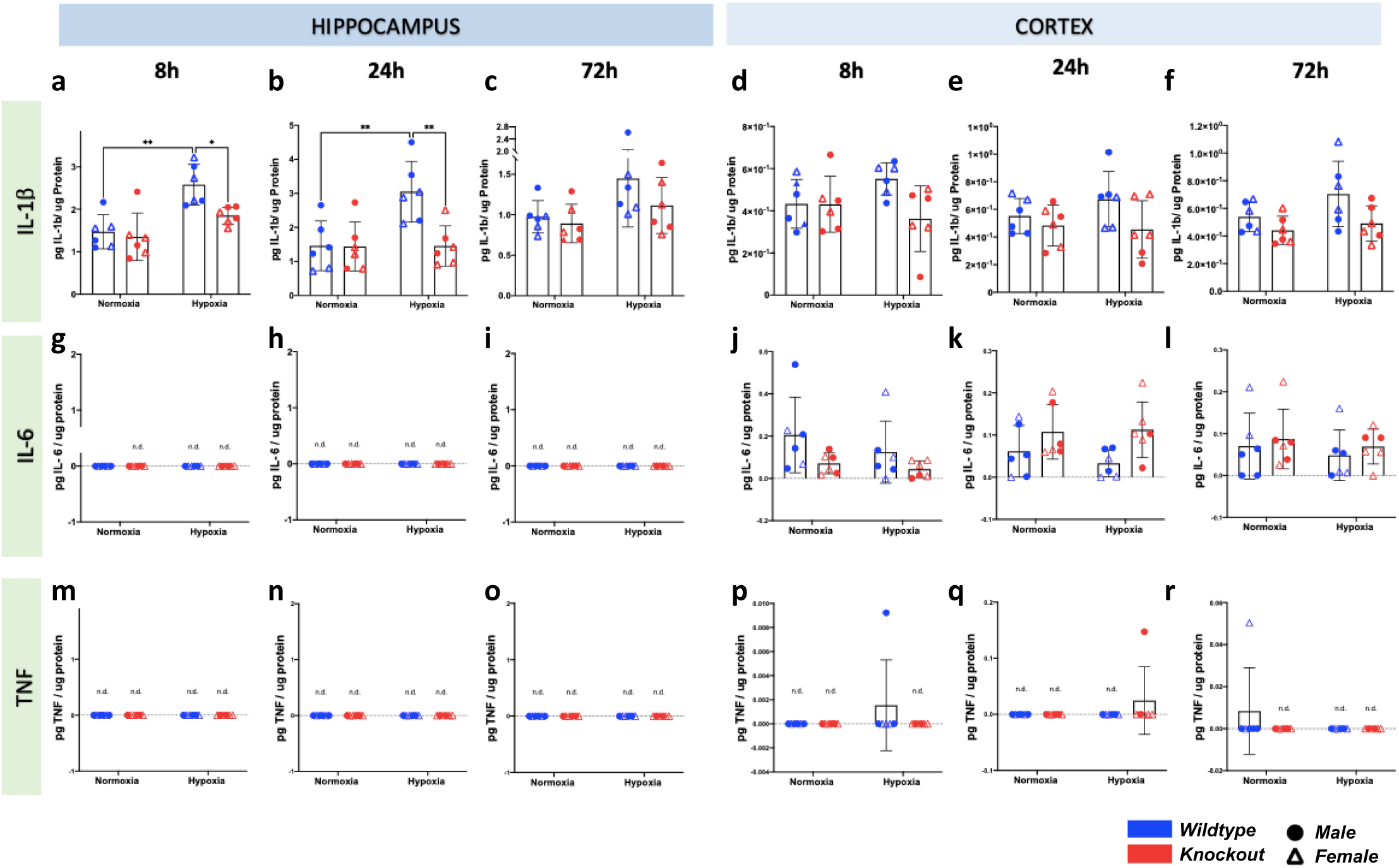
Modulation of inflammatory cytokines at the protein level in neonatal mice in response to globalised hypoxia. *Control and experimental genotype p7 pups were exposed to normoxia or hypoxia (5% oxygen) for 10 minutes at 34° C. Pups were monitored for 1 hour after induction of seizures. Brain tissue was extracted 8, 24 or 72 hours after induction of seizures by hypoxia*. **(a-f)** Hippocampal and cortical IL-1β protein levels in neonatal mice with hypoxic brain injury at 8, 24 and 72 hours following induction of seizure phenotype **(g-l)** Hippocampal and cortical IL-6 protein levels in neonatal mice with hypoxic brain injury at 8, 24 and 72 hours following induction of seizure phenotype **(m-r)** Hippocampal and cortical TNF protein levels in neonatal mice with hypoxic brain injury at 8, 24 and 72 hours following induction of seizure phenotype Tissue was added to 250ul of low stringency lysis buffer (LSLB) and homogenized. Tissue lysate IL-1β protein levels were measured using enzyme linked immunosorbent assay (ELISA). Tissue lysate IL-6 and TNF levels were measured using cytometric bead array (CBA). BCA assay was used to determine the protein concentration of each lysate, which was used to calculate the total protein in mg present in each sample. Cytokine expression is expressed relative to the amount of total protein in the sample (e.g.: pg IL-6/ mg protein). Relative IL-1β, IL-6 and TNF expression is mean ± SD of 6 biological replicates of each experimental category. * p˂0.05, ** p˂0.01, ***p˂0.001, two-way ANOVA. pg – picograms, n.d. – not detected. TNF – Tumour Necrosis Factor.

**Supplemental Figure 6.**
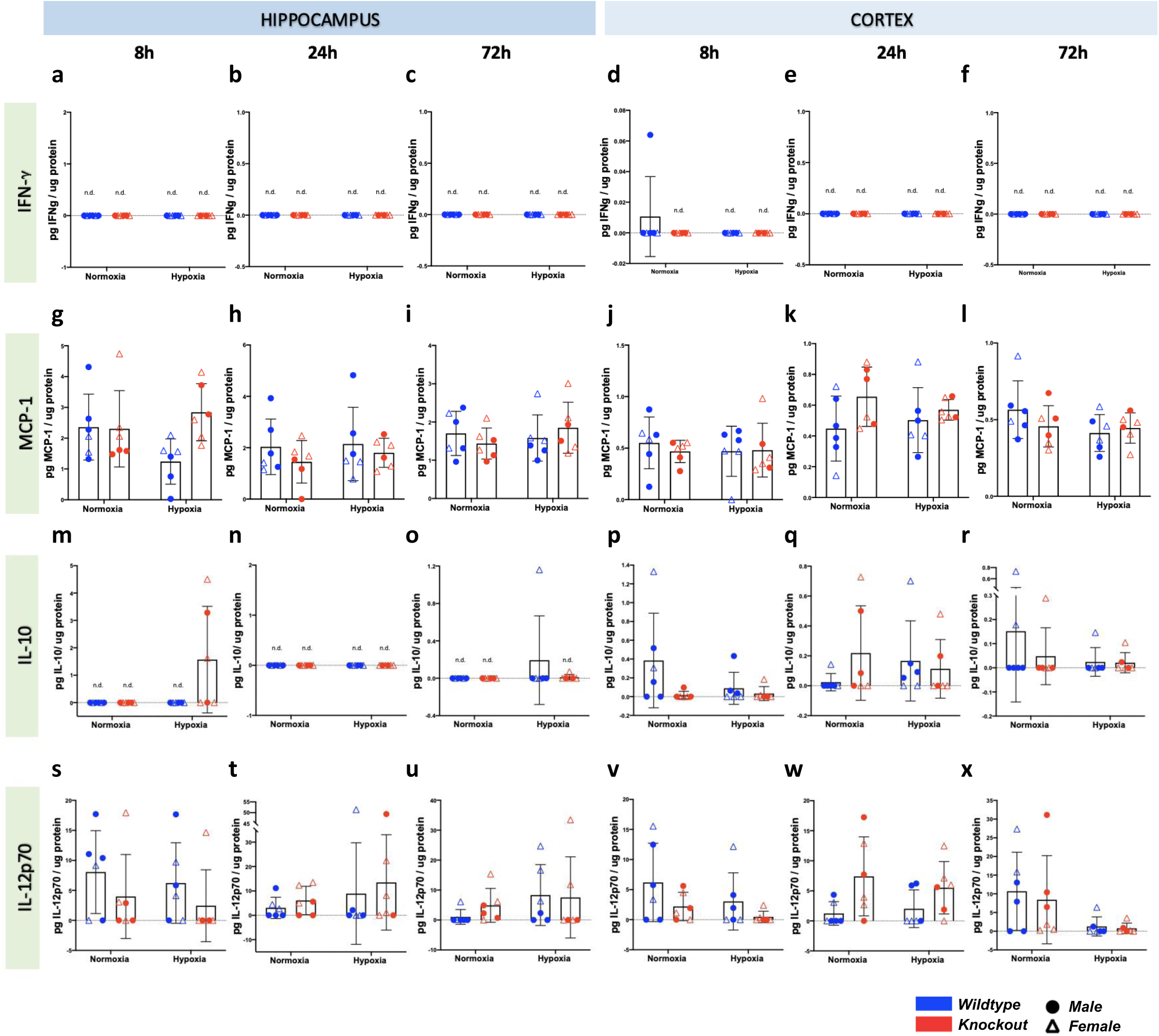
Modulation of inflammatory cytokines at the protein level in neonatal mice in response to globalised hypoxia. *Control and experimental genotype p7 pups were exposed to normoxia or hypoxia (5% oxygen) for 10 minutes at 34° C. Pups were monitored for 1 hour after induction of seizures. Brain tissue was extracted 8, 24 or 72 hours after induction of seizures by hypoxia*. **(a-f)** Hippocampal and cortical IFNγ protein levels in neonatal mice with hypoxic brain injury at 8, 24 and 72 hours following induction of seizure phenotype **(g-l)** Hippocampal and cortical MCP-1 protein levels in neonatal mice with hypoxic brain injury at 8, 24 and 72 hours following induction of seizure phenotype **(m-r)** Hippocampal and cortical IL-10 protein levels in neonatal mice with hypoxic brain injury at 8, 24 and 72 hours following induction of seizure phenotype **(s-x)** Hippocampal and cortical IL-12p70 protein levels in neonatal mice with hypoxic brain injury at 8, 24 and 72 hours following induction of seizure phenotype Tissue was added to 250ul of low stringency lysis buffer (LSLB) and homogenized. Tissue lysate cytokine levels were measured using cytometric bead array (CBA). BCA assay was used to determine the protein concentration of each lysate, which was used to calculate the total protein in mg present in each sample. Cytokine expression is expressed relative to the amount of total protein in the sample (e.g.: pg IL-6/ mg protein). Relative IFNγ, MCP-1, IL-10 and IL- 12p70 expression is mean ± SD of 6 biological replicates of each experimental category. * p˂0.05, ** p˂0.01, ***p˂0.001, two-way ANOVA. pg – picograms, n.d. – not detected.

**Supplemental Figure 7.**
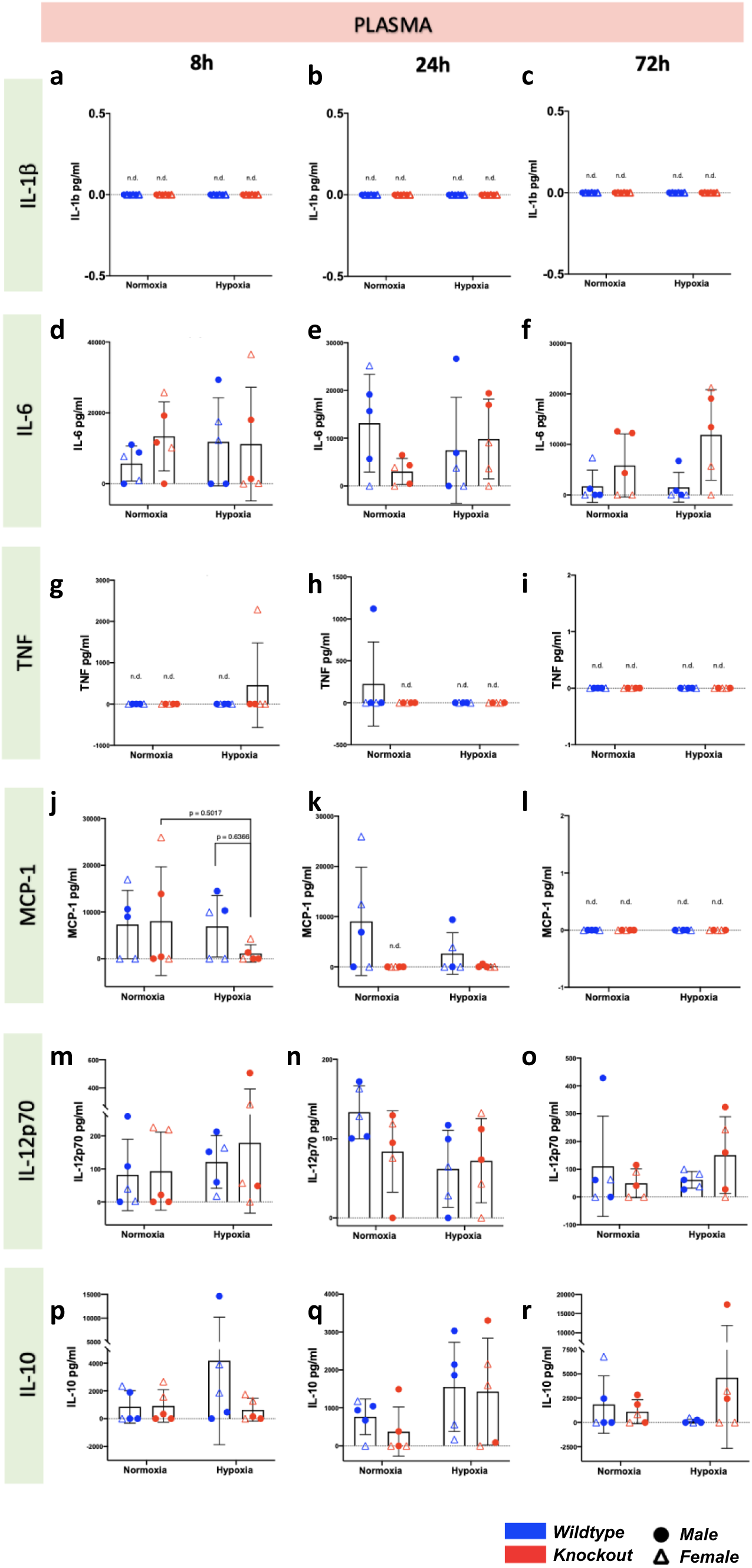
Plasma cytokine IL-1b (a-c), IL-6 (d-f), TNF (g-i) protein levels in p7 mice *in vivo*. Control and experimental genotype p7 pups were exposed to normoxia or hypoxia (5% oxygen) for 10 minutes at 34°C. Pups were monitored for 1 hour after induction of seizures. Blood was extracted 8, 24 or 72 hours after induction of seizures by hypoxia. **(a-c)** Serum IL-1β protein levels in neonatal mice with hypoxic brain injury at 8, 24 and 72 hours following induction of seizure phenotype **(d-f)** Serum IL-6 protein levels in neonatal mice with hypoxic brain injury at 8, 24 and 72 hours following induction of seizure phenotype **(g-i)** Serum TNF protein levels in neonatal mice with hypoxic brain injury at 8, 24 and 72 hours following induction of seizure phenotype **(j-l)** Serum MCP-1 protein levels in neonatal mice with hypoxic brain injury at 8, 24 and 72 hours following induction of seizure phenotype **(m-o)** Serum IL-12p70 protein levels in neonatal mice with hypoxic brain injury at 8, 24 and 72 hours following induction of seizure phenotype **(p-r)** Serum IL-10 protein levels in neonatal mice with hypoxic brain injury at 8, 24 and 72 hours following induction of seizure phenotype Blood was added to 15ml EDTA and centrifuged at 10,000rcf for 10 minutes to separate plasma. Plasma IL-1b protein levels were measured using enzyme linked immunosorbent assay (ELISA). Relative IL-1β expression is mean ± SD of 6 biological replicates of each experimental category. Plasma IL-6, TNF, IFNγ, MCP-1, IL-12p70 and IL-10 levels were measured using cytometric bead array. Relative IL-6, TNF, IFNγ, MCP-1, IL-12p70 and IL-10 expression is mean ± SD of 5 biological replicates of each experimental category.

**Supplemental Fig 8.**
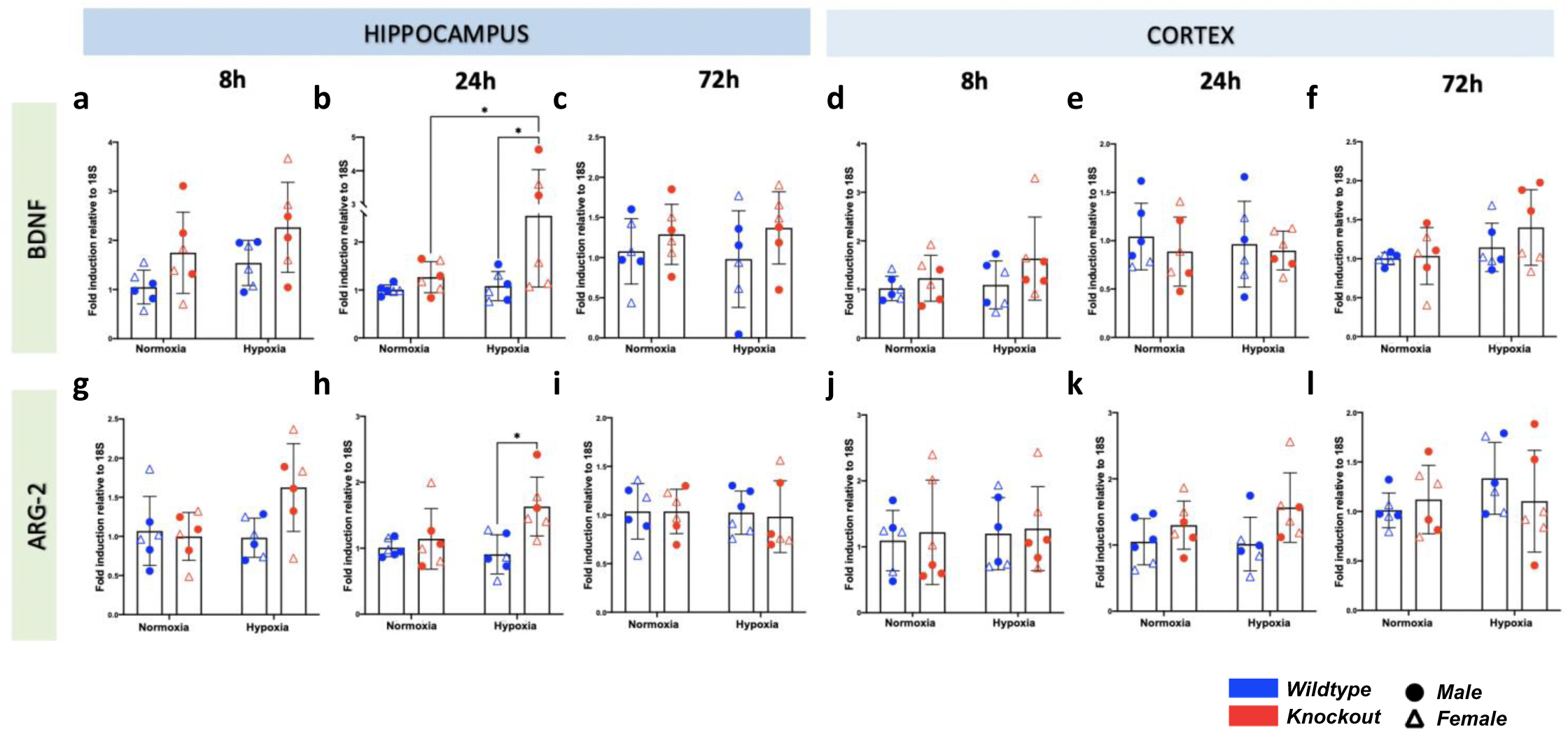
myeloid miR-155 deletion increases transcriptional expression of inflammatory regulators BDNF and ARG-2 in response to hypoxic brain injury. Control and experimental genotype p7 pups were exposed to normoxia or hypoxia (5% oxygen) for 10 minutes at 34°C. Pups were monitored for 1 hour after induction of seizures. Brain tissue was extracted 8, 24 or 72 hours after induction of seizures by hypoxia. **(a-f)** Hippocampal and cortical BDNF mRNA levels in neonatal mice with hypoxic brain injury at 8, 24 and 72 hours following induction of seizure phenotype **(g-l)** Hippocampal and cortical arg2 mRNA levels in neonatal mice with hypoxic brain injury at 8, 24 and 72 hours following induction of seizure phenotype RNA was extracted from tissue using TRIzol reagent protocols. cDNA was generated from samples and miR-155 or HIF-1α mRNA expression levels were evaluated using RT-qPCR. mRNA expression was normalised to 18s house-keeping gene and DCt of non-treated cells. Fold change was calculated using the formula 2-(DDCt). Relative expression is mean ± SD of 6 biological replicates of each experimental category. * p˂0.05, ** p˂0.01, ***p˂0.001, two- way ANOVA. BDNF – Brain Derived Neurotrophic Factor, ARG - Arginase

**Supplemental Fig 9.**
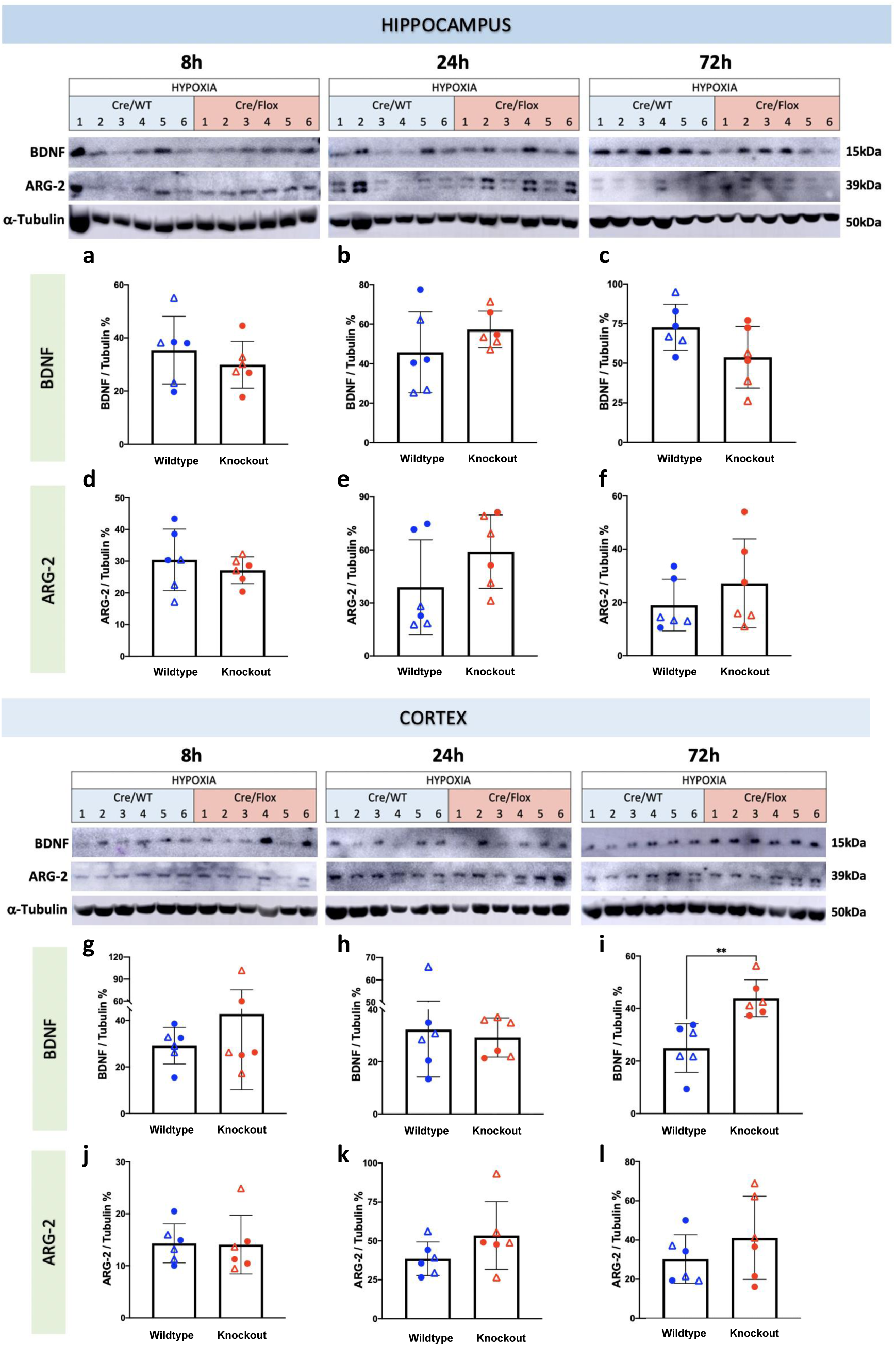
Protein levels of inflammatory regulators BDNF and ARG-2 in response to hypoxic brain injury. Control and experimental genotype p7 pups were exposed to normoxia or hypoxia (5% oxygen) for 10 minutes at 34°C. Pups were monitored for 1 hour after induction of seizures. Brain tissue was extracted 8, 24 or 72 hours after induction of seizures by hypoxia. **(a-c)** Hippocampal BDNF protein levels in neonatal mice with hypoxic brain injury at 8, 24 and 72 hours following induction of seizure phenotype **(d-f)** Hippocampal ARG-2 protein levels in neonatal mice with hypoxic brain injury at 8, 24 and 72 hours following induction of seizure phenotype **(g-i)** Cortical BDNF protein levels in neonatal mice with hypoxic brain injury at 8, 24 and 72 hours following induction of seizure phenotype **(j-l)** Cortical ARG-2 protein levels in neonatal mice with hypoxic brain injury at 8, 24 and 72 hours following induction of seizure phenotype Tissue was added to 250ul of low stringency lysis buffer (LSLB) and homogenized. BCA assay was used to determine the protein concentration of each lysate, which was used to calculate the volumes of lysate, dH20 and 5xSDS buffer to generate samples at 1000 mg/ml. Gel electrophoresis was performed on samples, and proteins were transferred onto polyvinylidene fluoride membranes. Membranes were blocked with 5% milk in TBS-T1x and coated with 1° and 2° antibodies. LumiGLO reagent and peroxide were added to membranes before performing Chemiluminescence. Relative BDNF and ARG-2 expression is mean ± SD of 6 biological replicates of each experimental category. BDNF – Brain Derived Neurotrophic Factor, ARG – Arginase

**Supplemental Fig 10.**
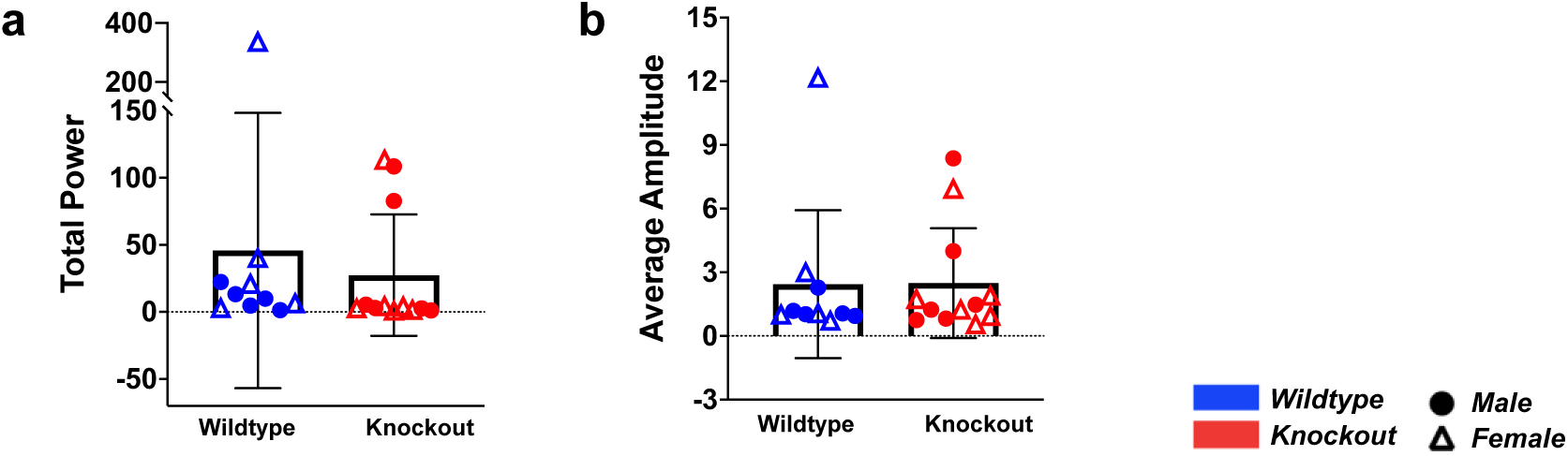
Myeloid miR-155 knockdown reduces seizure activity induced by hypoxic brain injury in the neonate. Electrodes were surgically implanted in control and experimental genotype p7 pups. Baseline recordings (minimum 20 minutes) of rest brain activity were taken from the mice after recovering from the surgery. These mice were then exposed hypoxia (5% oxygen) for 10 minutes at 34°C while continuous monitored using electroencephalography (EEG). Pups were monitored via EEG for 1 hour recovering after induction of seizures. **(a)** Total power of electroencephalographic outputs from neonatal mice during exposure to hypoxia **(b)** Average amplitude of electroencephalographic outputs from neonatal mice during exposure to hypoxia Power and amplitude during hypoxia and recovery were normalised to baseline read outs. Average power and amplitude ± SD were calculated from 7 biological replicates of the control genotype and 8 biological replicates of the conditional knockout experimental category (C and D). * p˂0.05, ** p˂0.01, ***p˂0.001, unpaired nonparametric t-test (Mann-Whitney test)(C), multiple t-tests (B and D).

## References

1. WHO. Children: Improving Survival And Well-Being. 2020 [30 October 2020]. Available from: https://www.who.int/news-room/fact-sheets/detail/children-reducing-mortality.

2. Fukuda S, Mizuno K, Kawai S, Kakita H, Goto T, Hussein MH, et al. Reduction in cerebral blood flow volume in infants complicated with hypoxic ischemic encephalopathy resulting in cerebral palsy. Brain and Development. 2008;30(4):246–53.

3. Eunson P. The long-term health, social, and financial burden of hypoxic–ischaemic encephalopathy. Developmental Medicine & Child Neurology. 2015;57(S3):48–50.

4. Ryan JM, Peterson MD, Matthews A, Ryan N, Smith KJ, O’Connell NE, et al. Noncommunicable disease among adults with cerebral palsy: A matched cohort study. Neurology. 2019;93(14):e1385–e96.

5. Smith KJ, Peterson MD, O’Connell NE, Victor C, Liverani S, Anokye N, et al. Risk of Depression and Anxiety in Adults With Cerebral Palsy. JAMA Neurology. 2019;76(3):294–300.

6. Shankaran S, Pappas A, McDonald SA, Vohr BR, Hintz SR, Yolton K, et al. Childhood outcomes after hypothermia for neonatal encephalopathy. N Engl J Med. 2012;366(22):2085–92.

7. Guillet R, Edwards AD, Thoresen M, Ferriero DM, Gluckman PD, Whitelaw A, et al. Seven- to eight-year follow-up of the CoolCap trial of head cooling for neonatal encephalopathy. Pediatr Res. 2012;71(2):205–9.

8. Pisani F, Orsini M, Braibanti S, Copioli C, Sisti L, Turco EC. Development of epilepsy in newborns with moderate hypoxic-ischemic encephalopathy and neonatal seizures. Brain Dev. 2009;31(1):64–8.

9. Kossoff E. Neonatal seizures due to hypoxic-ischemic encephalopathy: should we care? Epilepsy Curr. 2011;11(5):147–8.

10. Kharoshankaya L, Stevenson NJ, Livingstone V, Murray DM, Murphy BP, Ahearne CE, et al. Seizure burden and neurodevelopmental outcome in neonates with hypoxic–ischemic encephalopathy. Developmental Medicine & Child Neurology. 2016;58(12):1242–8.

11. Miller SM, Goasdoue K, Björkman ST. Neonatal seizures and disruption to neurotransmitter systems. Neural Regen Res. 2017;12(2):216–7.

12. Soul JS. Acute symptomatic seizures in term neonates: Etiologies and treatments. Semin Fetal Neonatal Med. 2018;23(3):183–90.

13. Bhalala US, Koehler RC, Kannan S. Neuroinflammation and neuroimmune dysregulation after acute hypoxic-ischemic injury of developing brain. Front Pediatr. 2015;2:144-.

14. Rocha-Ferreira E, Hristova M. Plasticity in the Neonatal Brain following Hypoxic-Ischaemic Injury. Neural Plast. 2016;2016:4901014-.

15. Davidson JO, Wassink G, van den Heuij LG, Bennet L, Gunn AJ. Therapeutic Hypothermia for Neonatal Hypoxic-Ischemic Encephalopathy - Where to from Here? Front Neurol. 2015;6:198-.

16. Perlman JM. Pathogenesis of hypoxic-ischemic brain injury. Journal of Perinatology. 2007;27(1):S39–S46.

17. Tsitsiou E, Lindsay MA. microRNAs and the immune response. Curr Opin Pharmacol. 2009;9(4):514–20.

18. Guarnieri DJ, DiLeone RJ. MicroRNAs: A new class of gene regulators. Annals of Medicine. 2008;40(3):197–208.

19. Carissimi C, Fulci V, Macino G. MicroRNAs: Novel regulators of immunity. Autoimmunity Reviews. 2009;8(6):520–4.

20. Momen-Heravi F, Bala S. miRNA regulation of innate immunity. Journal of Leukocyte Biology. 2018;103(6):1205–17.

21. Alivernini S, Gremese E, McSharry C, Tolusso B, Ferraccioli G, McInnes IB, et al. MicroRNA-155—at the Critical Interface of Innate and Adaptive Immunity in Arthritis. Frontiers in Immunology. 2018;8(1932).

22. Bruning U, Cerone L, Neufeld Z, Fitzpatrick SF, Cheong A, Scholz CC, et al. MicroRNA- 155 promotes resolution of hypoxia-inducible factor 1alpha activity during prolonged hypoxia. Mol Cell Biol. 2011;31(19):4087–96.

23. Wan G, Xie W, Liu Z, Xu W, Lao Y, Huang N, et al. Hypoxia-induced MIR155 is a potent autophagy inducer by targeting multiple players in the MTOR pathway. Autophagy. 2014;10(1):70–9.

24. Wan G, Xie W, Liu Z, Xu W, Lao Y, Huang N, et al. Hypoxia-induced MIR155 is a potent autophagy inducer by targeting multiple players in the MTOR pathway. Autophagy. 2014;10(1):70–9.

25. Chan SY, Loscalzo J. MicroRNA-210: a unique and pleiotropic hypoxamir. Cell Cycle. 2010;9(6):1072–83.

26. Caballero-Garrido E, Pena-Philippides JC, Lordkipanidze T, Bragin D, Yang Y, Erhardt EB, et al. In Vivo Inhibition of miR-155 Promotes Recovery after Experimental Mouse Stroke. J Neurosci. 2015;35(36):12446–64.

27. Pena-Philippides JC, Caballero-Garrido E, Lordkipanidze T, Roitbak T. In vivo inhibition of miR-155 significantly alters post-stroke inflammatory response. J Neuroinflammation. 2016;13(1):287-.

28. Xing G, Luo Z, Zhong C, Pan X, Xu X. Influence of miR-155 on Cell Apoptosis in Rats with Ischemic Stroke: Role of the Ras Homolog Enriched in Brain (Rheb)/mTOR Pathway. Med Sci Monit. 2016;22:5141–53.

29. Roitbak T. Silencing a Multifunctional microRNA Is Beneficial for Stroke Recovery. Front Mol Neurosci. 2018;11:58-.

30. Zhang L, Liu C, Huang C, Xu X, Teng J. miR-155 Knockdown Protects against Cerebral Ischemia and Reperfusion Injury by Targeting MafB. BioMed Research International. 2020;2020:6458204.

31. Stoll G, Jander S, Schroeter M. Inflammation and glial responses in ischemic brain lesions. Prog Neurobiol. 1998;56(2):149–71.

32. Wang X, Yue T-L, Barone FC, Feuerstein GZ. Monocyte Chemoattractant Protein&#x2013;1 Messenger RNA Expression in Rat Ischemic Cortex. Stroke. 1995;26(4):661–6.

33. Kim JS, Gautam SC, Chopp M, Zaloga C, Jones ML, Ward PA, et al. Expression of monocyte chemoattractant protein-1 and macrophage inflammatory protein-1 after focal cerebral ischemia in the rat. Journal of Neuroimmunology. 1995;56(2):127–34.

34. Murdoch C, Muthana M, Lewis CE. Hypoxia regulates macrophage functions in inflammation. J Immunol. 2005;175(10):6257–63.

35. Fang H-Y, Hughes R, Murdoch C, Coffelt SB, Biswas SK, Harris AL, et al. Hypoxia-inducible factors 1 and 2 are important transcriptional effectors in primary macrophages experiencing hypoxia. Blood. 2009;114(4):844–59.

36. Liu F, McCullough LD. Inflammatory responses in hypoxic ischemic encephalopathy. Acta Pharmacologica Sinica. 2013;34(9):1121–30.

37. Albina JE, Henry WL, Jr., Mastrofrancesco B, Martin BA, Reichner JS. Macrophage activation by culture in an anoxic environment. J Immunol. 1995;155(9):4391–6.

38. Scannell G. Leukocyte responses to hypoxic/ischemic conditions. New Horiz. 1996;4(2):179–83.

39. Murata Y, Ohteki T, Koyasu S, Hamuro J. IFN-gamma and pro-inflammatory cytokine production by antigen-presenting cells is dictated by intracellular thiol redox status regulated by oxygen tension. Eur J Immunol. 2002;32(10):2866–73.

40. Jablonski KA, Gaudet AD, Amici SA, Popovich PG, Guerau-de-Arellano M. Control of the Inflammatory Macrophage Transcriptional Signature by miR-155. PLoS One. 2016;11(7):e0159724-e.

41. Sly LM, Rauh MJ, Kalesnikoff J, Song CH, Krystal G. LPS-induced upregulation of SHIP is essential for endotoxin tolerance. Immunity. 2004;21(2):227–39.

42. Costinean S, Sandhu SK, Pedersen IM, Tili E, Trotta R, Perrotti D, et al. Src homology 2 domain-containing inositol-5-phosphatase and CCAAT enhancer-binding protein beta are targeted by miR-155 in B cells of Emicro-MiR-155 transgenic mice. Blood. 2009;114(7):1374–82.

43. O’Connell RM, Chaudhuri AA, Rao DS, Baltimore D. Inositol phosphatase SHIP1 is a primary target of miR-155. Proc Natl Acad Sci U S A. 2009;106(17):7113–8.

44. McCoy CE, Sheedy FJ, Qualls JE, Doyle SL, Quinn SR, Murray PJ, et al. IL-10 inhibits miR-155 induction by toll-like receptors. J Biol Chem. 2010;285(27):20492–8.

45. O’Neill LA, Sheedy FJ, McCoy CE. MicroRNAs: the fine-tuners of Toll-like receptor signalling. Nature Reviews Immunology. 2011;11(3):163–75.

46. Cardoso AL, Guedes JR, Pereira de Almeida L, Pedroso de Lima MC. miR-155 modulates microglia-mediated immune response by down-regulating SOCS-1 and promoting cytokine and nitric oxide production. Immunology. 2012;135(1):73–88.

47. O’Connell RM, Rao DS, Baltimore D. microRNA Regulation of Inflammatory Responses. Annual Review of Immunology. 2012;30(1):295–312.

48. Self-Fordham JB, Naqvi AR, Uttamani JR, Kulkarni V, Nares S. MicroRNA: Dynamic Regulators of Macrophage Polarization and Plasticity. Front Immunol. 2017;8:1062-.

49. Vigorito E, Kohlhaas S, Lu D, Leyland R. miR-155: an ancient regulator of the immune system. Immunological Reviews. 2013;253(1):146–57.

50. Jing W, Zhang X, Sun W, Hou X, Yao Z, Zhu Y. CRISPR/CAS9-Mediated Genome Editing of miRNA-155 Inhibits Proinflammatory Cytokine Production by RAW264.7 Cells. Biomed Res Int. 2015;2015:326042.

51. Mann M, Mehta A, Zhao JL, Lee K, Marinov GK, Garcia-Flores Y, et al. An NF-κB- microRNA regulatory network tunes macrophage inflammatory responses. Nat Commun. 2017;8(1):851.

52. Androulidaki A, Iliopoulos D, Arranz A, Doxaki C, Schworer S, Zacharioudaki V, et al. The kinase Akt1 controls macrophage response to lipopolysaccharide by regulating microRNAs. Immunity. 2009;31(2):220–31.

53. Lu LF, Thai TH, Calado DP, Chaudhry A, Kubo M, Tanaka K, et al. Foxp3-dependent microRNA155 confers competitive fitness to regulatory T cells by targeting SOCS1 protein. Immunity. 2009;30(1):80–91.

54. Wang P, Hou J, Lin L, Wang C, Liu X, Li D, et al. Inducible microRNA-155 Feedback Promotes Type I IFN Signaling in Antiviral Innate Immunity by Targeting Suppressor of Cytokine Signaling 1. The Journal of Immunology. 2010;185(10):6226–33.

55. Wang C, Zhang C, Liu L, A X, Chen B, Li Y, et al. Macrophage-Derived mir-155- Containing Exosomes Suppress Fibroblast Proliferation and Promote Fibroblast Inflammation during Cardiac Injury. Mol Ther. 2017;25(1):192–204.

56. Zhang Z, Wang Z, Zhang B, Liu Y. Downregulation of microRNA-155 by preoperative administration of valproic acid prevents postoperative seizures by upregulating SCN1A. Mol Med Rep. 2018;17(1):1375–81.

57. Ashhab MU, Omran A, Kong H, Gan N, He F, Peng J, et al. Expressions of tumor necrosis factor alpha and microRNA-155 in immature rat model of status epilepticus and children with mesial temporal lobe epilepsy. J Mol Neurosci. 2013;51(3):950–8.

58. Huang L-G, Zou J, Lu Q-C. Silencing rno-miR-155-5p in rat temporal lobe epilepsy model reduces pathophysiological features and cell apoptosis by activating Sestrin-3. Brain Research. 2018;1689:109–22.

59. Cai Z, Li S, Li S, Song F, Zhang Z, Qi G, et al. Antagonist Targeting microRNA-155 Protects against Lithium-Pilocarpine-Induced Status Epilepticus in C57BL/6 Mice by Activating Brain-Derived Neurotrophic Factor. Front Pharmacol. 2016;7:129-.

60. Zhang W, Wang L, Pang X, Zhang J, Guan Y. Role of microRNA-155 in modifying neuroinflammation and γ-aminobutyric acid transporters in specific central regions after post-ischaemic seizures. Journal of Cellular and Molecular Medicine. 2019;23(8):5017–24.

61. Rodriguez-Alvarez N, Jimenez-Mateos EM, Dunleavy M, Waddington JL, Boylan GB, Henshall DC. Effects of hypoxia-induced neonatal seizures on acute hippocampal injury and later-life seizure susceptibility and anxiety-related behavior in mice. Neurobiology of Disease. 2015;83:100–14.

62. Quinlan S, Merino-Serrais P, Di Grande A, Dussmann H, Prehn JHM, Ní Chonghaile T, et al. The Anti-inflammatory Compound Candesartan Cilexetil Improves Neurological Outcomes in a Mouse Model of Neonatal Hypoxia. Frontiers in Immunology. 2019;10(1752).

63. Lagos-Quintana M, Rauhut R, Yalcin A, Meyer J, Lendeckel W, Tuschl T. Identification of Tissue-Specific MicroRNAs from Mouse. Current Biology. 2002;12(9):735–9.

64. Jenkins DD, Rollins LG, Perkel JK, Wagner CL, Katikaneni LP, Bass WT, et al. Serum cytokines in a clinical trial of hypothermia for neonatal hypoxic-ischemic encephalopathy. J Cereb Blood Flow Metab. 2012;32(10):1888–96.

65. Dowling JK, Afzal R, Gearing LJ, Cervantes-Silva MP, Annett S, Davis GM, et al. Mitochondrial arginase-2 is essential for IL-10 metabolic reprogramming of inflammatory macrophages. Nature Communications. 2021;12(1):1460.

66. Quinlan S, Merino-Serrais P, Di Grande A, Dussmann H, Prehn JHM, Ni Chonghaile T, et al. The Anti-inflammatory Compound Candesartan Cilexetil Improves Neurological Outcomes in a Mouse Model of Neonatal Hypoxia. Front Immunol. 2019;10:1752.

67. Soul JS. Acute symptomatic seizures in term neonates: Etiologies and treatments. Seminars in Fetal and Neonatal Medicine. 2018;23(3):183–90.

68. Glass HC, Shellhaas RA, Wusthoff CJ, Chang T, Abend NS, Chu CJ, et al. Contemporary Profile of Seizures in Neonates: A Prospective Cohort Study. The Journal of Pediatrics. 2016;174:98–103.e1.

69. Nair J, Kumar VHS. Current and Emerging Therapies in the Management of Hypoxic Ischemic Encephalopathy in Neonates. Children (Basel). 2018;5(7):99.

70. Oorschot DE, Sizemore RJ, Amer AR. Treatment of Neonatal Hypoxic-Ischemic Encephalopathy with Erythropoietin Alone, and Erythropoietin Combined with Hypothermia: History, Current Status, and Future Research. Int J Mol Sci. 2020;21(4):1487.

71. Allen KA, Brandon DH. Hypoxic Ischemic Encephalopathy: Pathophysiology and Experimental Treatments. Newborn Infant Nurs Rev. 2011;11(3):125–33.

72. Brennan GP, Henshall DC. MicroRNAs as regulators of brain function and targets for treatment of epilepsy. Nature Reviews Neurology. 2020;16(9):506–19.

73. Reschke CR, Henshall DC. microRNA and Epilepsy. Adv Exp Med Biol. 2015;888:41–70.

74. Henshall DC. MicroRNA and epilepsy: profiling, functions and potential clinical applications. Curr Opin Neurol. 2014;27(2):199–205.

75. Duan W, Chen Y, Wang XR. MicroRNA-155 contributes to the occurrence of epilepsy through the PI3K/Akt/mTOR signaling pathway. Int J Mol Med. 2018;42(3):1577–84.

76. Fu H, Cheng Y, Luo H, Rong Z, Li Y, Lu P, et al. Silencing MicroRNA-155 Attenuates Kainic Acid-Induced Seizure by Inhibiting Microglia Activation. Neuroimmunomodulation. 2019;26(2):67–76.

77. Wang J, Zhao J. MicroRNA Dysregulation in Epilepsy: From Pathogenetic Involvement to Diagnostic Biomarker and Therapeutic Agent Development. Frontiers in Molecular Neuroscience. 2021;14(35).

78. Cai Z, Li S, Li S, Song F, Zhang Z, Qi G, et al. Antagonist Targeting microRNA-155 Protects against Lithium-Pilocarpine-Induced Status Epilepticus in C57BL/6 Mice by Activating Brain-Derived Neurotrophic Factor. Frontiers in Pharmacology. 2016;7(129).

79. Aly H, Khashaba MT, El-Ayouty M, El-Sayed O, Hasanein BM. IL-1beta, IL-6 and TNF-alpha and outcomes of neonatal hypoxic ischemic encephalopathy. Brain Dev. 2006;28(3):178–82.

80. Bajnok A, Berta L, Orbán C, Veres G, Zádori D, Barta H, et al. Distinct cytokine patterns may regulate the severity of neonatal asphyxia—an observational study. J Neuroinflammation. 2017;14(1):244.

81. Marathe C, Bradley MN, Hong C, Lopez F, Ruiz de Galarreta CM, Tontonoz P, et al. The arginase II gene is an anti-inflammatory target of liver X receptor in macrophages. J Biol Chem. 2006;281(43):32197–206.

82. Krystofova J, Pathipati P, Russ J, Sheldon A, Ferriero D. The Arginase Pathway in Neonatal Brain Hypoxia-Ischemia. Dev Neurosci. 2018;40(5-6):437–50.

83. Yin Y, Phạm TL, Shin J, Shin N, Kang D-W, Lee SY, et al. Arginase 2 Deficiency Promotes Neuroinflammation and Pain Behaviors Following Nerve Injury in Mice. J Clin Med. 2020;9(2):305.

84. Caballero-Garrido E, Pena-Philippides JC, Lordkipanidze T, Bragin D, Yang Y, Erhardt EB, et al. <em>In Vivo</em> Inhibition of miR-155 Promotes Recovery after Experimental Mouse Stroke. The Journal of Neuroscience. 2015;35(36):12446–64.

85. Pena-Philippides JC, Caballero-Garrido E, Lordkipanidze T, Roitbak T. In vivo inhibition of miR-155 significantly alters post-stroke inflammatory response. J Neuroinflammation. 2016;13(1):287.

86. Hsin J-P, Lu Y, Loeb GB, Leslie CS, Rudensky AY. The effect of cellular context on miR- 155-mediated gene regulation in four major immune cell types. Nature Immunology. 2018;19(10):1137–45.

87. O’Connell RM, Taganov KD, Boldin MP, Cheng G, Baltimore D. MicroRNA-155 is induced during the macrophage inflammatory response. Proceedings of the National Academy of Sciences. 2007;104(5):1604–9.

88. Li Q, Barres BA. Microglia and macrophages in brain homeostasis and disease. Nature Reviews Immunology. 2018;18(4):225–42.

89. Bachiller S, Jiménez-Ferrer I, Paulus A, Yang Y, Swanberg M, Deierborg T, et al. Microglia in Neurological Diseases: A Road Map to Brain-Disease Dependent-Inflammatory Response. Frontiers in Cellular Neuroscience. 2018;12(488).

90. Andoh M, Ikegaya Y, Koyama R. Synaptic Pruning by Microglia in Epilepsy. J Clin Med. 2019;8(12):2170.

91. Hill CA, Fitch RH. Sex Differences in Mechanisms and Outcome of Neonatal Hypoxia-Ischemia in Rodent Models: Implications for Sex-Specific Neuroprotection in Clinical Neonatal Practice. Neurology Research International. 2012;2012:867531.

92. Netto CA, Sanches E, Odorcyk FK, Duran-Carabali LE, Weis SN. Sex-dependent consequences of neonatal brain hypoxia-ischemia in the rat. Journal of Neuroscience Research. 2017;95(1-2):409–21.

93. Chan CB, Ye K. Sex differences in brain-derived neurotrophic factor signaling and functions. Journal of neuroscience research. 2017;95(1-2):328–35.

94. Varendi K, Kumar A, Härma MA, Andressoo JO. miR-1, miR-10b, miR-155, and miR- 191 are novel regulators of BDNF. Cell Mol Life Sci. 2014;71(22):4443–56.

95. Solich J, Kuśmider M, Faron-Górecka A, Pabian P, Kolasa M, Zemła B, et al. Serum Level of miR-1 and miR-155 as Potential Biomarkers of Stress-Resilience of NET-KO and SWR/J Mice. Cells. 2020;9(4):917.

96. Falcicchia C, Paolone G, Emerich DF, Lovisari F, Bell WJ, Fradet T, et al. Seizure- Suppressant and Neuroprotective Effects of Encapsulated BDNF-Producing Cells in a Rat Model of Temporal Lobe Epilepsy. Molecular Therapy - Methods & Clinical Development. 2018;9:211–24.

97. Binder DK, Croll SD, Gall CM, Scharfman HE. BDNF and epilepsy: too much of a good thing? Trends Neurosci. 2001;24(1):47–53.

98. Scharfman HE. Brain-derived neurotrophic factor and epilepsy--a missing link? Epilepsy Curr. 2005;5(3):83–8.

99. Augusto-Oliveira M, Arrifano GP, Lopes-Araújo A, Santos-Sacramento L, Takeda PY, Anthony DC, et al. What Do Microglia Really Do in Healthy Adult Brain? Cells. 2019;8(10):1293.

